# Deficiency of the Heterogeneous Nuclear Ribonucleoprotein U locus leads to delayed hindbrain neurogenesis

**DOI:** 10.1101/2022.09.14.507275

**Authors:** Francesca Mastropasqua, Marika Oksanen, Cristina Soldini, Shemim Alatar, Abishek Arora, Roberto Ballarino, Maya Molinari, Federico Agostini, Axel Poulet, Michelle Watts, Ielyzaveta Rabkina, Martin Becker, Danyang Li, Britt-Marie Anderlid, Johan Isaksson, Karl Lundin Remnelius, Mohsen Moslem, Yannick Jacob, Anna Falk, Nicola Crosetto, Magda Bienko, Emanuela Santini, Anders Borgkvist, Sven Bölte, Kristiina Tammimies

## Abstract

Genetic variants affecting *Heterogeneous Nuclear Ribonucleoprotein U (HNRNPU)* have been identified in several neurodevelopmental disorders (NDDs). HNRNPU is widely expressed in the human brain and shows the highest postnatal expression in the cerebellum. Recent studies have investigated the role of *HNRNPU* in cerebral cortical development, but the effects of *HNRNPU* deficiency on cerebellar development remain unknown. Here, we describe the molecular and cellular outcomes of *HNRNPU* locus deficiency during *in vitro* neural differentiation of patient-derived and isogenic neuroepithelial stem cells with a hindbrain profile. We demonstrate that *HNRNPU* deficiency leads to chromatin remodeling of A/B compartments, and transcriptional rewiring, partly by impacting exon inclusion during mRNA processing. Genomic regions affected by the chromatin restructuring and host genes of exon usage differences show a strong enrichment for genes implicated in epilepsies, intellectual disability, and autism. Lastly, we show that at the cellular level. *HNRNPU* downregulation leads to altered neurogenesis and an increased fraction of neural progenitors in the maturing neuronal population. We conclude that, *HNRNPU* locus is involved in delayed commitment of neural progenitors to neuronal maturation in cell types with hindbrain profile.

## INTRODUCTION

Enormous progress in genomic technologies has led to the discovery of hundreds of genes associated with various neurodevelopmental disorders (NDDs). These include several genes belonging to the heterogeneous nuclear ribonucleoprotein (hnRNP) family (Gillentine et al., 2021). One of the genes within this family, *HNRNPU*, which encodes for Heterogeneous Nuclear Ribonucleoprotein U (also known as Scaffold Attachment Factor A (SAF-A))(Kiledjian and Dreyfuss, 1992), has emerged as a frequently affected gene leading to NDDs such as intellectual disability (ID), autism spectrum disorder (ASD) as well as neurological conditions such as epilepsies (Depienne et al., 2017; Satterstrom et al., 2020; Taylor et al., 2022). The gene was first indicated in NDDs as part of the 1q44 microdeletion syndrome characterized by a severe global developmental delay, ID, seizures, muscular hypotonia, hearth, and congenital malformations such as agenesis of the corpus callosum, heart and skeletal anomalies(Depienne et al., 2017). Later, several smaller deletions and point mutations affecting the *HNRNPU* locus pinpointed it as the causal gene within the locus for the majority of the brain related phenotypes(Bramswig et al., 2017; Leduc et al., 2017; Shimojima et al., 2012; Tung et al., 2021; Yates et al., 2017). In addition to the *HNRNPU* gene, a long non-coding RNA *HNRNPU-AS1* maps to the locus. The function of *HNRNPU-AS1* is not known, although some reports have indicated its role in different molecular pathways, such as cell proliferation and apoptosis in cancer cells (Niu et al., 2021; Zhang et al., 2022). To date, several reports exist about the pathogenic genetic variants affecting the *HNRNPU* gene in individuals with *HNRNPU*-related disorder, and are mostly *de novo* loss-of-function variants at sequence or copy number level (Brunet et al., 2021; Durkin et al., 2020; Gillentine et al., 2021; Wang et al., 2020). All the reported cases are heterozygous variants, suggesting that homozygous gene-disrupting variants affecting *HNRNPU* are embryonic lethal in humans, similar to what has been demonstrated in mice (Dugger et al., 2020; Roshon and Ruley, 2005; Ye et al., 2015).

HNRNPU has a key role in 3D genome organization and regulating RNA processing (Fan et al., 2018; Marenda et al., 2022; Nozawa et al., 2017; Xiong et al., 2020; Ye et al., 2015). For instance, HNRNPU modulates chromatin compaction by promoting chromatin accessibility in a dynamic yet structured fashion, dependent on its oligomerization status (Nozawa et al., 2017). Recent studies have also shown the involvement of HNRNPU in mitosis and cell division by changing its interactions with condensed chromatin and influencing DNA replication and sister chromatid separation (Connolly et al., 2022; Ma et al., 2011; Sharp et al., 2020). Several studies have reported a role for HNRNPU in splicing, promoting both exon inclusion and exclusion (Huelga et al., 2012; Jones et al., 2022; Ye et al., 2015). Mechanistically, it has been shown that HNRNPU can stabilize the pre-mRNA structure, thus inhibiting the splicing of certain exons (Jones et al., 2022). A critical role for Hnrnpu-mediated splicing has also been demonstrated during pre- and postnatal heart development in mice, showing that loss of *Hnrnpu* leads to increased intron retention events, causing abnormality in heart development and function (Ye et al., 2015).

Recently, few studies shed light on the effects of *HNRNPU* mutations in cortical development. A mouse model of *Hnrnpu* haploinsufficiency presented abnormal brain organization and showed at postnatal day 0 altered transcriptome with downregulation of pathways related to neuronal projection and migration and upregulation of genes relevant to cell growth and protein localization in hippocampal and neocortical cells (Dugger et al., 2020). In contrast, a study performed on embryonic mice upon complete conditional *Hnrnpu* knockout in the cerebral cortex showed upregulation of genes involved in synaptic activity and downregulation of DNA-related ontologies together with changes in alternative splicing regulation (Sapir et al., 2022). Similarly, an isogenic human cortex organoid model with two clonal cell lines carrying different heterozygous mutations in *HNRNPU* partially resembled what was observed in the embryonic mice but not the postnatal model (Ressler et al., 2023). Therefore, the overall outcome of *HNRNPU* genetic variants might depend on the stage of development, brain region studied, genetic background, and gene dosage. So far, the studies have mainly focused on cortex development and forebrain structures. *HNRNPU* is expressed in different tissues, and postnatally it is highest in cerebellum (Thierry et al., 2012). Accordingly, atrophy of the cerebellum was highlighted in a cohort of patients affected by *HNRNPU*-related disorder (Durkin et al., 2020). The cerebellum is one of the most studied hindbrain structures, and has emerged as important for typical and atypical development, and abnormalities in cerebellar development have been associated with ASD and ID (Burstein and Geva, 2021; Frosch et al., 2022; Spahiu et al., 2022). Recent observations have highlighted the influence of hindbrain development on the brain cortex and its pivotal role for cognitive functions (Joseph, R., n.d.; Kohlmeier and Polli, 2020) and suggested that analysis of midbrain and hindbrain are needed to expand the knowledge about NDDs (London et al., 2022).

Here, we focused on delineating the molecular and cellular consequences of the *HNRNPU* locus deficiency in a model of human early hindbrain development using induced pluripotent stem cells (iPSCs) from an individual with *HNRNPU*-related disorder and a knockdown isogenic cell approach for comparative analyses, and providing evidence of a broad spectrum of affected pathways, converging towards delayed neurogenesis.

## RESULTS

### Generation and characterization of *HNRNPU* locus knockdown in iPS and neuroepithelial stem cells

To assess the molecular effects of *HNRNPU* locus (including *HNRNPU* and *HNRNPU-AS1*) haploinsufficiency during human hindbrain differentiation, we generated two different *HNRNPU* locus deficient cellular models derived from human iPSCs. The iPSCs were induced into neuroepithelial stem (NES) cells with hindbrain profile and further differentiated for 5 (D5) and 28 (D28) days using an undirected protocol as previously described (Becker et al., 2020; Falk et al., 2012) (**Figure 1A-B**). This approach generates a neuronal culture consisting of a mixed population of excitatory, inhibitory, and progenitor cells representing a physiological neuronal environment (Falk et al., 2012).

**Figure 1.**
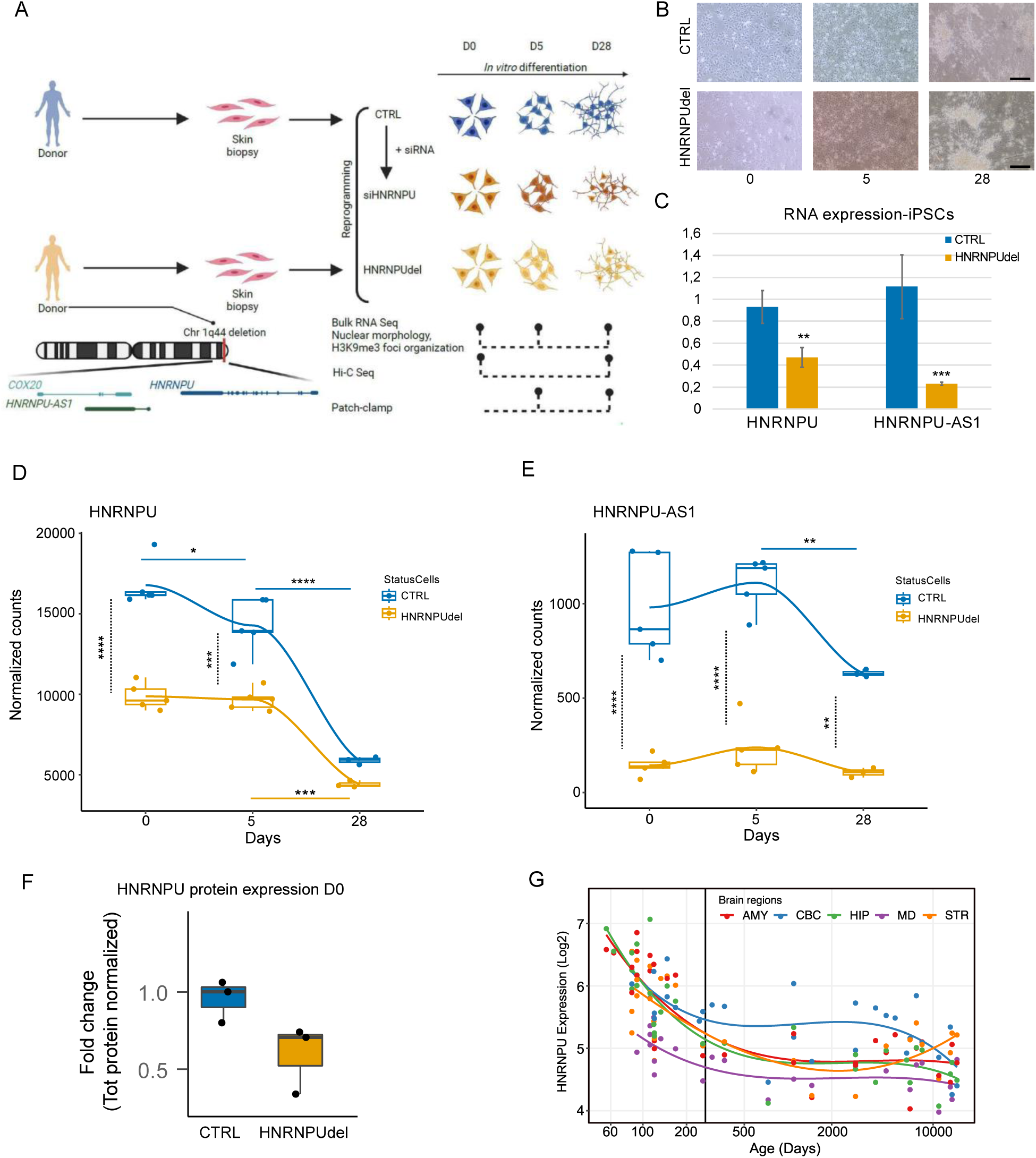
*HNRNPU* expression changes during neural differentiation and brain development. **A)** Schematic summary of the samples and methods used in the study. **B)** Brightfield microscopy of CTRL and HNRNPU_del/+_ cells at D0, D5, D28. Scale bar= 50 µm. **C)** *HNRNPU* and *HNRNPU-AS1* RNA expression in CTRL and HNRNPU_del/+_ iPSC cells. **D-E)** *HNRNPU* (**D**) and *HNRNPU-AS1*(**E**) RNA expression in CTRL and HNRNPU_del/+_ at D0, D5, D28. The full lines indicate the comparisons between the time points for each cell line; the black dotted line indicates the comparison between the two cell lines at each time point. **F)** HNRNPU protein expression in CTRL and HNRNPU_del/+_ at D0. **G)** *HNRNPU* RNA expression in all the brain regions during development from Human Brain Transcriptome dataset. The vertical line indicates the time of birth. AMY= Amygdala, CBC= Cerebellar Cortex, HIP= Hippocampus, MD= Mediodorsal nucleus thalamus, STR= Striatum \**p*<0.05; \*\**p*<1×10^-3^; \*\*\**p*<1×10^-4^; \*\*\*\**p*<1×10^-5^. In the figure, the HNRNPU_del/+_ samples are indicated as “HNRNPUdel”.

We first generated an *HNRNPU*-related disorder patient cell model, hereafter called HNRNPU_del/+_. Through genetic screening of a twin cohort focusing on NDDs, we identified a male twin pair carrying a 44 kilobase (kb) heterozygous deletion spanning from *COX20* to *HNRNPU* genes (Stamouli et al., 2018). The twins were diagnosed with ASD, ID, and fever-induced seizures. A detailed phenotypic description of the twin pair is presented in **Supplementary Table 1**. We obtained fibroblasts from skin biopsies of the twins and successfully reprogrammed twin-2 fibroblasts into iPSCs. The iPSCs had a normal karyotype, pluripotent marker expression and showed significant reduction of *HNRNPU* and *HNRNPU-AS1* compared with control iPSCs (**Figure 1C and Figure S1A, S1B**). In addition, we used dual-SMAD inhibition to derive NES cells from the iPSCs as previously described (Falk et al., 2012), followed by staining for neuronal stem cell markers Nestin and SRY (sex determining region Y)-box 2 (SOX2) to verify their identity (**Figure S1C**). To confirm the downregulation of *HNRNPU*-related RNA and protein product, we measured *HNRNPU-AS1* RNA, and *HNRNPU* RNA and protein in HNRNPU_del/+_ cells using CTRL cells as a reference. HNRNPU_del/+_ showed significantly lower RNA expression for both *HNRNPU-AS1* and *HNRNPU* (*P* < 1.0×10^−5^, ANOVA and posthoc Tukey test) and an average lower expression of HNRNPU protein spanning from ∼20% to ∼60% downregulation (**Figure 1D-F**).

As a complementary approach, we generated an isogenic cell model, hereafter called siHNRNPU, in which we reduced *HNRNPU-AS1* RNA, and *HNRNPU* RNA and protein expression using a pool of small interfering RNA (siRNA) oligos in NES cells obtained from a neurotypical male control (CTRL) (Uhlin et al., 2017a) in parallel with non-target oligo pool (siNTC), similar to approaches successfully used in previous studies of HNRNPU in other cell types (Ma et al., 2011; Nozawa et al., 2017; Zietzer et al., n.d.) . To achieve consistent knockdown of *HNRNPU* throughout differentiation, we performed repetitive siRNA treatments every six days. Significant downregulation of *HNRNPU-AS1* and *HNRNPU* RNA and a 32% reduction of HNRNPU protein expression were observed at NES stage. Both transcripts were similarly significantly downregulated at D5, but no significant difference was observed in *HNRNPU* RNA expression after 28 days in differentiation either by RNA sequencing (RNA-seq) or real-time PCR (**Figure S1D**). Nonetheless, HNRNPU protein expression was reduced to 29% at D28 compared with siNTC samples (**Figure S1G**), therefore, we considered the silencing successful and proceeded with further analyses. The *HNRNPU-AS1* was consistently downregulated after differentiation (**Figure S1D**).

Our results of the variable RNA and protein expressions after downregulation of *HNRNPU* by siRNA treatment or mutation are consistent with all the previous studies about *HNRNPU* mutations in brain tissues and differentiated neuronal populations (Dugger et al., 2020; Ressler et al., 2023; Sapir et al., 2022). Even after knockdown or knockout at gene level, these studies showed that *HNRNPU* expression levels are similar to the wild type controls suggesting possible compensatory mechanisms. A possible mechanism might reside in the capacity of proteins belonging to the HNRNP family to directly associate with their own transcripts, thus stabilizing them (Huelga et al., 2012). For instance, single-cell RNA transcriptome comparison of human cortical organoids carrying two different frameshift mutations in *HNRNPU* revealed no difference in HNRNPU expression between the HNRNPU-mutant and relative control, in any of the identified cell types. At the same time, the downregulation of *HNRNPU* at the RNA level does not always translate into a similar reduction at the protein level (Dugger et al., 2020; Ressler et al., 2023; Sapir et al., 2022).

### *HNRNPU* expression changes during hindbrain neural differentiation and brain development

Next, we sought to analyze the molecular consequences of *HNRNPU* haploinsufficiency during human hindbrain development. To this end, we extracted total RNA for transcriptomic analyses from NES cells collected at three different time points of differentiation (D0, D5, and D28). To confirm that our model resembles human hindbrain development, we analyzed the expression of several hindbrain and cerebellar markers in our cell line at NES and D28. We first analyzed the expression of *HOXA2* and *OTX1/OTX2* in neural stem cell/progenitor phases in our cell model as the balance of these markers is fundamental for the specification of early hindbrain development (Lowenstein et al., n.d.). At D0 and D5, the cells expressed *HOXA2* and not *OTX1/OTX2*, in line with the developing hindbrain phenotype (**Figure S1H**). Accordingly, we show that CTRL cells at D28 express many cerebellar markers (*UNC5C, ICMT, CA8, TRPC3, ASTN1, KITLG*) (Aldinger et al., 2021; Consalez et al., 2021; Kim et al., 2003; Kim and Ackerman, 2011; Mancarci et al., 2017; Wu et al., 2019)(**Figure S1I**). In parallel, we analyzed cerebellar marker expression in our previously published single-cell RNA-seq (scRNAseq) data from a similarly derived cell line at D28(Becker et al., 2020) and showed that the cerebellar markers are indeed expressed across the cell types (**Figure S1J**). Furthermore, to evaluate the specificity of our model, we analyzed the expression of the markers in a previously published dataset from human cortical organoids(Ressler et al., 2023) and observed extremely low or null expression of the markers in all the cell types and samples (**Figure S1K**).

When analyzing the expression of the *HNRNPU* and *HNRNPU-AS1* in the CTRL cell line, we observed that *HNRNPU-AS1* expression decreases from D5 to D28 (comparison D0-D5: *P* = 0.76; D5-D28: *P* < 0.005, ANOVA and posthoc Tukey test), and *HNRNPU* expression decreased steadily during differentiation (comparison D0-D5: *P* = 0.019; D5-D28: *P* = < 1.0×10^−5^, ANOVA and posthoc Tukey test) (**Figure 1D-E**). *HNRNPU* expression follows a similar decreasing trend during differentiation from iPSCs to neurons from a previously published dataset(Burke et al., 2020) and during development of all the cerebral areas, although in the cerebellum the postnatal expression is the highest compared to the other brain regions (**Figure 1G, Figure S1L).** Since *HNRNPU* expression was higher in proliferating progenitor cells (**Figure 1D** and (Ressler et al., 2023; Sapir et al., 2022)), we further inspected its expression in our above mentioned scRNA-seq data (**Figure S2A**). As expected, *HNRNPU* expression was highest in the neural progenitor population (**Figure S2B-C**). Moreover, the neural progenitor population was further divided into three distinct subclusters, of which one was highly enriched in proliferating markers such as *TOP2A*, *KIFC1,* and *KIF18B.* This cluster also had the highest expression of *HNRNPU* (**Figure S2D**).

In contrast to CTRL cells the HNRNPU_del/+_ cells did not show a change in the expression of *HNRNPU-AS1* during differentiation (comparison D0-D5: *P* = 0.92; D5-D28: *P* =0.84, ANOVA and posthoc Tukey test) (**Figure 1E**). Similarly, *HNRNPU* expression was stable from D0 to D5 (*P* = 0.99, ANOVA and posthoc Tukey test), but followed a significantly delayed decrease at D28 (D5-D28, *P* = 2.6×10^−5^, ANOVA and posthoc Tukey test) (**Figure 1D**). In the comparison of RNA expression between CTRL and HNRNPU_del/+_ at each time point, *HNRNPU-AS1* was significantly reduced in HNRNPU_del/+_ at each time point (D0 and D5: *P* < 1.0×10^−5^; D28: *P* =0.005, ANOVA and posthoc Tukey test) (**Figure 1E**). Instead, at D0 and D5, when the cells express progenitor markers Nestin and SOX2 (**Figure S3A**), *HNRNPU* expression was significantly lower in HNRNPU_del/+_ cells compared with CTRL cells (*P* < 1.0×10^−5^ and 2.3 ×10^−5^, respectively, ANOVA and posthoc Tukey test), while the expression was similar between the two cell lines at D28 (*P* = 0.58, ANOVA and posthoc Tukey test) (**Figure 1D**). HNRNPU protein was predominantly localized in the nucleus at all three analyzed stages of differentiation, and the measured protein levels were extremely variable between different time points in both cell lines and did not mirror mRNA levels (**Figure S3B-C**). These results show that physiologically *HNRNPU* has the highest expression in the neuroepithelial stem cell stage and steadily reduces during the differentiation demonstrating its important role in early neural differentiation.

### *HNRNPU* locus expression impacts cell differentiation pathways

Next, we performed differential gene expression analyses using DESeq2(Love et al., 2014), followed by gene set enrichment analysis (GSEA) of the obtained transcriptomic data. Similar to earlier reported results in other cell types (Nozawa et al., 2017), we found that *HNRNPU* downregulation had a limited effect on the transcriptional landscape at D0 and D5. The isogenic siHNRNPU cells (replicates n=5) had only 10 differentially expressed genes (DEG) at D0 and 30 DEG at D5 (Base Mean > 20, |log2FoldChange| > 0.58, *P* adjusted < 0.05, Wald test and Benjamini-Hochberg procedure) (**Supplementary Table 2A-B**). As expected, due to the different genetic backgrounds, when comparing HNRNPU_del/+_ cells (replicates n=5) to the CTRL cell line (replicates n=5) at each time point, we identified a higher number of DEG (2091 DEG at D0 of which 1033 upregulated and 1058 downregulated; 2091 DEG at D5, of which 1251 upregulated and 840 downregulated genes) (**Supplementary Table 2C-D**). At D28 of the neural differentiation, both *HNRNPU*-deficient models revealed wider transcriptional rewiring, with 1511 DEG in siHNRNPU and 1608 DEG in HNRNPU_del/+_ cells (**Supplementary Table 2E-F**). Of these, only 148 DEG genes were shared between the two models at D28, suggesting that downregulation of *HNRNPU* might affect the expression of upstream transcriptional regulators that increase the transcriptomic landscape variability.

Next, we analyzed gene set and pathway level changes across the two datasets (**Supplementary Table 3**). At D0, no specific Gene Ontology (GO) terms or enriched pathways were significantly shared between the two models (**Figure 2A, Figure S4A, and Supplementary Table 3A-D**). At D5, genes affecting the positive regulation of excitatory postsynaptic potential were downregulated in both models. In contrast, among upregulated genes, shared GO terms and enriched pathways included categories referring to the regulation of cell differentiation, growth factor receptors, and constituents of extracellular matrix (**Figure 2B, Figure S4B, and Supplementary Table 3E-H**).

**Figure 2.**
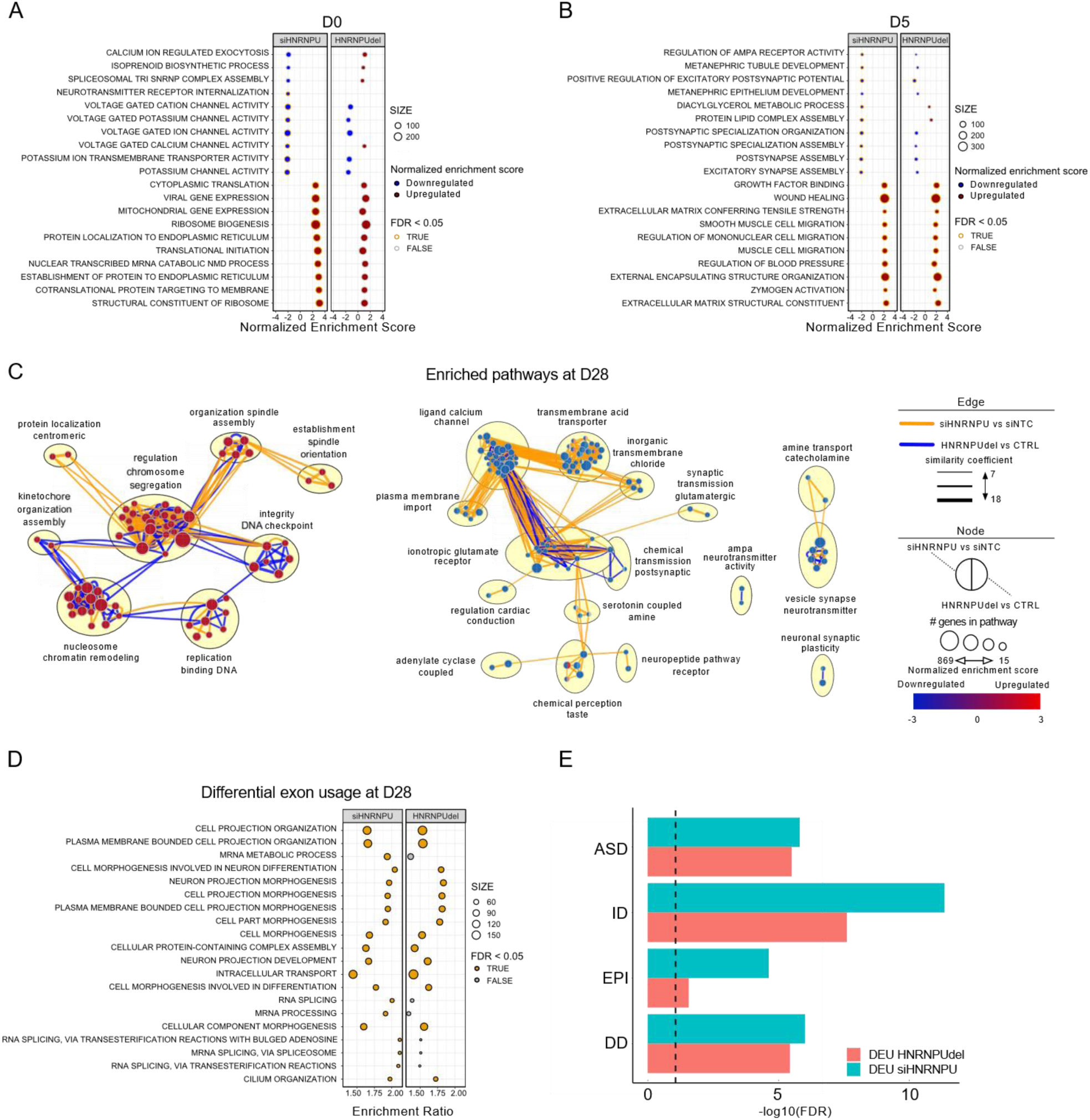
*HNRNPU* expression affects cell differentiation pathways and modulates exon usage of NDD genes. **A-B)** Top 10 upregulated and downregulated GO terms from ranked differentially expressed genes in siHNRNPU (left panel) and HNRNPU_del/+_ (right panel) at D0 (A) and D5 (B). **C)** Selected upregulated (left panel) and downregulated (right panel) pathways enriched at D28. For each node, the left half indicates the enrichment in siHNRNPU versus siNTC, and the right half the enrichment in HNRNPU_del/+_ versus CTRL. The color of the edge indicates which of the datasets significantly contributed to the pathway call (p adj <0.05). **D)** Top 20 GO terms enriched from genes with differential exon usage in siHNRNPU (left panel) and HNRNPU_del/+_ (right panel) at D28. **E)** Enrichment of genes subject to differential exon usage (DEU) in siHNRNPU versus siNTC (cyan) or HNRNPU_del/+_ versus CTRL (magenta) in ASD, ID, epilepsy (EPI), DD gene lists at D28. The vertical dotted line represents the significance threshold of FDR<0.05 (-log10(FDR)>1.3) after hypergeometric analysis. In the figure, the HNRNPU_del/+_ samples are indicated as “HNRNPUdel”.

The major transcriptional changes at D28 were clustering in multiple biological processes affected in both HNRNPU_del/+_ and siHNRNPU cells. We found 74 significantly enriched GO terms shared between the two models, including several synaptic and transmembrane channel ontologies among the downregulated pathways (**Supplementary Table 3I-L**). Interestingly, the shared upregulated pathways from GSEAs included pathways related to cell DNA organization during DNA replication and cell division and more general nucleosome and chromatin remodeling pathways (**Figure 2D**), confirming the role of HNRNPU in the regulation of chromatin organization and DNA replication even in neural cells. Moreover, we found that two key developmental pathways, epithelium tube and embryonic hindlimb morphogenesis, were significantly upregulated in both model systems. Additional dysregulated developmental pathways were also uniquely affected in HNRNPU_del/+_ (**Figure S4C**). We also investigated whether the DEG genes at D28 were enriched in genes previously associated with epilepsies, ID, ASD, and general developmental disorders (DD). However, we found no significant enrichment for any gene lists (Hypergeometric test, **Supplementary Table 4**). These results demonstrate that *HNRNPU* deficiency modulates the transcriptomic variability by altering the expression of genes enriched in neural maturation and chromatin organization and mostly affecting cells to be committed to neuronal differentiation more than cells in the neural progenitor phase.

### HNRNPU modulates exon usage of NDD genes

Since HNRNPU has previously been shown to affect the transcriptome by regulating alternative splicing (Sapir et al., 2022; Xiao et al., 2012; Ye et al., 2015), we analyzed differential exon usage (DEU), which indicates alternative splicing events or differential isoform usage. Similar to the gene level changes, we detected fewer DEU events (ExonBaseMean > 10, |Log2FoldChange| > 0.58 and P adjusted < 0.05) in the earlier timepoints and more at D28 in both model systems (**Figure S4D and Supplementary Table 5**). When comparing siHNRNPU to siNTC samples, we detected 0 and 2 DEU genes at D0 and D5, respectively, whereas a comparison between HNRNPU_del/+_ and CTRL yielded 35 and 38 DEU genes at D0 and D5, respectively. At D28, 976 and 1285 DEU genes were detected for siHNRNPU and HNRNPU_del/+_ when compared against their respective controls, respectively. Notably, only 5.9% and 9.8% of DEU genes were also differentially expressed in siHNRNPU and HNRNPU_del/+_ cells, respectively. Both models revealed more exclusion than inclusion of exons due to *HNRNPU* deficiency, as 71% of the DEU events were due to downregulation (**Figure S4D**). This result is in line with previous findings using other cell types (Huelga et al., 2012; Sapir et al., 2022; Ye et al., 2015). Over representation analysis (ORA) of the DEU genes shared by both models identified enriched pathways involved in cell morphogenesis, neuron projection development, and cilium organization (**Figure 2D and Supplementary Table 6**). We also analyzed whether DEU genes were enriched for the genes implicated in the different disorders as earlier and found a strong enrichment for ID gene list (hypergeometric test followed with false discovery rate (FDR) correction: 1.49×10^−14^ and 5.26×10^−10^ for siHNRNPU and HNRNPU_del/+_, respectively), ASD (8.26×10^−8^ and 1.96×10^−7^, respectively), DD (4.68×10^−8^ and 2.36×10^−7^), and epilepsy (2.28×10^−6^ and 0.0121) (**Figure 2E and Supplementary Table 4**). Our results demonstrate that HNRNPU is involved in regulating exon usage during neural development, of genes previously associated with several NDDs.

### *HNRNPU* locus deficiency increases the proportion of neural progenitor cells during differentiation

Transcriptional changes at the pathway level strongly indicated differences in the cell proliferation rate of HNRNPU_del/+_ and siHNRNPU cells at D28; therefore, we focused on analyzing the neural progenitor pool at D28, a timepoint in which generally most of the cells are postmitotic and committed for neuronal maturation. First, we analyzed the cell type proportions using deconvolution of the transcriptomic data similar to previously described (Becker et al., 2020). The deconvolution predicted a higher proportion of neural progenitors in HNRNPU_del/+_ compared to CTRL (**Figure 3A and Figure S5A**). To validate the presence of neural progenitors across the differentiation and investigate the difference between the two models, we analyzed SOX2 positive nuclei at the three-time points for both HNRNPU_del/+_ and CTRL cell lines. As expected, each cell line showed a decreasing number of SOX2-positive cells during the differentiation time course. However, HNRNPU_del/+_ cells displayed a significantly higher number of SOX2-positive cells at D5 and D28 compared to CTRL (*P* < 10^−4^ and 0.002, respectively, ꭓ^2^ test) (**Figure 3B and Figure S5B**). Also, the siHNRNPU model had an increased number of SOX2-positive cells at D28, but the difference was insignificant (*P* = 0.16, ꭓ^2^ test) compared with siNTC cells (**Figure 3B**). To evaluate the characteristics of the progenitor cells at D28, we measured cell proliferation by Bromo-deoxy-uridine (BrdU) incorporation in both conditions. As expected, the cell proliferation rate decreased throughout the differentiation and was significantly higher in both siHNRNPU and HNRNPU_del/+_ cells at D28 compared to the relative controls at the same time point (*P* = 0.002 and 0.01, respectively, Two-sided T-test). The rate was similar between HNRNPU_del/+_ and CTRL cell lines at D0 but diverged already at D5 (*P* = 6×10^−3^, Two-sided T-test) (**Figure 3C and Figure S5C**). Altogether this demonstrates that upon *HNRNPU* locus downregulation, the mixed population at D28 is enriched in proliferating neural progenitor cells compared with the control conditions, suggesting a delayed maturation trajectory during differentiation.

**Figure 3.**
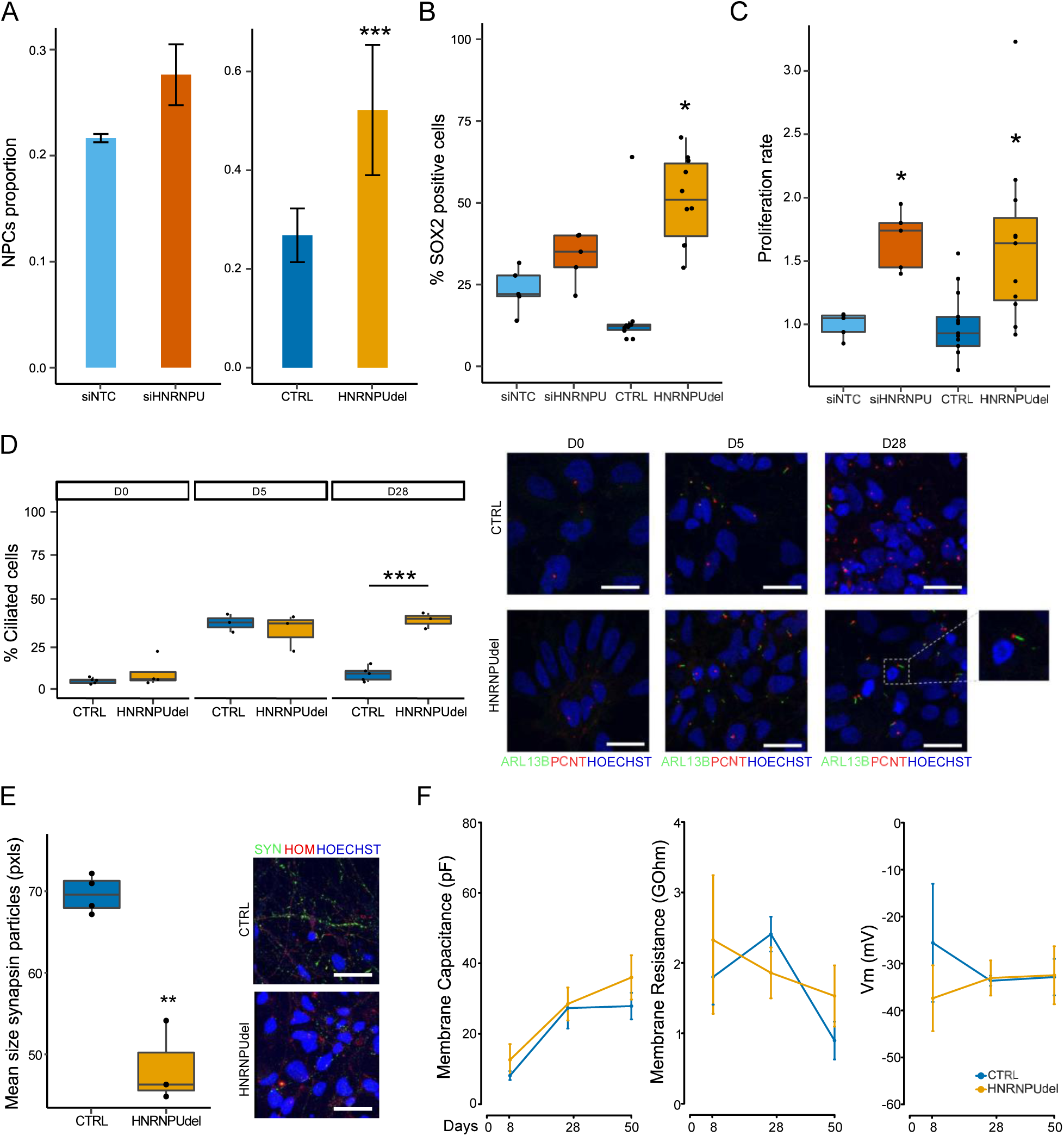
Cells at the late-differentiation stage show higher progenitor phenotype and affected synaptogenesis under *HNRNPU* downregulation. **A)** Neural progenitor cells proportion in siNTC, siHNRNPU, CTRL, HNRNPU_del/+_ at D28 estimated by deconvolution analysis. **B)** Percentage of cells positive at the staining with SOX2 antibody at D28. **C)** Proliferation rate of siNTC, siHNRNPU, CTRL, HNRNPU_del/+_ at D28. **D)** Immunofluorescence of primary cilia with ciliary marker ARL13B and basal body marker PCNT in CTRL and HNRNPU_del/+_ and quantification of ciliated cell proportions. Pictures were acquired with 63X magnification, 0.5 zoom and z-stack. Scale bar= 20 µm. **E)** Immunofluorescence of synapsin1/2 in CTRL and HNRNPU_del/+_ at D28 and quantification of the mean size of the synapse particles positive to synapsin1/2 staining. Pictures were acquired with 63X magnification and z-stack. Scale bar= 20 µm. **F)** Electric properties of the membrane of CTRL and HNRNPU_del/+_ cells at different time points. \**p* < 0.05, \*\**p* < 0.001, *** < 0.0001. In the figure HNRNPU_del/+_ samples are indicated as “HNRNPUdel”.

### *HNRNPU* locus downregulation alters the maturation of neural cells

Since the dysregulated pathways in both *HNRNPU*-deficient conditions were related to membrane channels, synaptic formation, and extracellular matrix, we hypothesized that *HNRNPU* deficiency could affect the stage of neuronal maturation. To validate this hypothesis and the transcriptomic results, we first analyzed the proportion of cells with primary cilia during neural differentiation, as primary cilia guide axon tract development (Guo et al., 2019). Additionally, our results for DEU genes showed the enrichment of cilium organization in siHNRNPU and HNRNPUdel/+ cells at D28. Therefore, we analyzed ciliary proteins ARL13 and PCNT expression by immunofluorescence in CTRL and HNRNPU_del/+_ cells throughout differentiation and observed a higher number of ciliated cells at D5 compared to D0 in both cell lines and no difference between *HNRNPU* deficiency and control cells at D0 and D5 (P=0.37 and P=0.64, respectively, ꭓ^2^-test). However, a significant difference in the proportion of ciliated cells was observed at D28 (P=1.497×10^-6^, ꭓ^2^-test), when HNRNPU_del/+_ cells had a significantly higher percentage of ciliated cells (**Figure 3D**). Since the alteration of cilium organization pathways was similarly significant in both experimental conditions, we considered sufficient to validate the finding by immunofluorescence only in the HNRNPU_del/+_ condition.

Additionally, we analyzed the expression of the presynaptic marker Synapsin 1/2 and the postsynaptic marker Homer1. The HNRNPU_del/+_ cells had significantly smaller presynaptic particle sizes than the CTRL cells (*P* = 0.009, T-test, two-sided) (**Figure 3E**). In contrast, the number of synaptic particles and the number and size of postsynaptic signals were comparable between HNRNPU_del/+_ and CTRL. To monitor the neuronal maturation, we performed patch clamp electrophysiology to study the intrinsic membrane properties and synaptic activity of HNRNPU_del/+_ and CTRL cells at D8 (n = 4), D28 (n = 12 cells for CTRL, and 17 cells for HNRNPU_del/+_), and D50 (n = 7 cells for CTRL and 10 cells for HNRNPU_del/+_) (**Figure 3F**). However, despite the clear neuronal morphology (**Figure S5D**), the cells were characterized by high membrane resistance in response to voltage-step commands, and no spontaneous excitatory or inhibitory synaptic currents could be measured in any of the recorded cells, therefore at this stage of differentiation neither CTRL nor HNRNPU_del/+_ cells can be considered mature neurons. These results indicate that the cilia-guided neuronal maturation process is delayed in cells with *HNRNPU* deficiency.

### *HNRNPU* locus downregulation affects nuclear shape and chromatin organization

The results of our transcriptional profiling revealed a role of HNRNPU in chromatin organization, in line with previous reports that demonstrated the function of HNRNPU in chromatin compaction, DNA synthesis, and chromosome folding during mitosis in different cell types and conditions (Connolly et al., 2022; Nozawa et al., 2017; Sharp et al., 2020). Therefore, we sought to investigate whether *HNRNPU* deficiency affects chromatin organization in our human neural cell model. First, we performed chromatin accessibility analyses by treating CTRL and HNRNPU_del/+_ cells with DNaseI; however, no large-scale differences were visible (**Figure S6A**). We then performed single-cell immunofluorescent analysis of cell nuclei stained for heterochromatin marker, histone 3 tri-methylation at lysine 9 (H3K9me3). Both *HNRNPU*-deficient cell models displayed profound changes in nuclear architecture at D28 (**Figure 4A**). Specifically, upon *HNRNPU* deficiency, a subpopulation of cells displayed differential number and total volume of heterochromatic foci (NbF, VFTotal, respectively), intensity and volume of the relative heterochromatic fraction (Intensity RHF, Volume RHF), and nuclear shape characteristics like surface area, volume, and radius of a sphere of equivalent volume (ESR) (**Figure 4A**).

**Figure 4.**
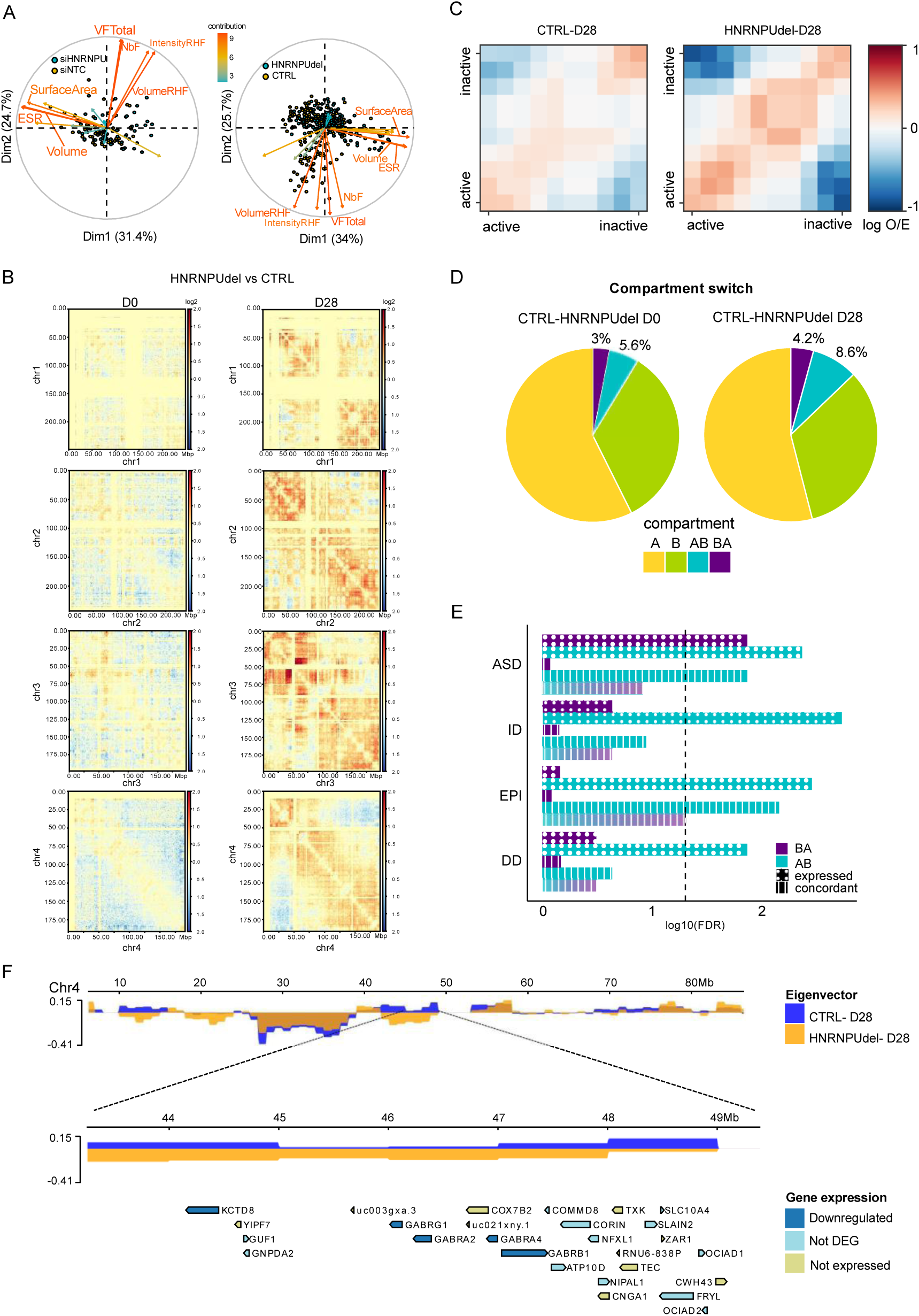
*HNRNPU* downregulation affects chromatin organization. **A)** Biplot of the contribution of the variables generated by NucleusJ for the clustering of the nuclei after H3K9me3 staining, compared siHNRNPU with siNTC (left panel) and HNRNPU_del/+_ with CTRL (right panel). **B)** Example of normalized log2ratio of the contacts for each chromosome of HNRNPU _del/+_ compared with CTRL, at D0 and D28, after ICE correction. **C)** Saddle plot of the *cis-* and *trans-* interactions of the A (active) and B (inactive) compartments in CTRL and HNRNPU_del/+_ at D28. **D)** Genome-wise compartment switch at D0 and D28 in the comparison for each time point of HNRNPU_del/+_ and CTRL. **E)** An enrichment of genes within the compartment switch regions in ASD, ID, epilepsy (EPI), and DD gene lists at D28. “Expressed”, indicated by the dotted pattern, include genes that map in one of the compartment switches and are expressed in our dataset (Gene count > 20 in at least one sample). “Concordant”, indicated by the vertical lines pattern, includes the genes that are upregulated from the transcriptome analysis and map in a compartment B in CTRL and A in HNRNPU_del/+_ or vice versa. The vertical dotted black line represents the significance threshold of FDR<0.05 (-log10(FDR)>1.3) after the hypergeometric test corrected for all the comparisons. **F)** Representative compartment switch from A to B at D28 between CTRL and HNRNPU_del/+_ on chromosome 4. The upper panel shows the eigenvectors of the first 90 Mb of chr 4 for each sample (yellow for HNRNPU_del/+_ and blue for CTRL). The zoom-in panel shows the eigenvectors of the two samples in the chromosome location of chr4: 43-49 Mb. The lowest panel shows the genes mapping onto the switch region (Dark blue: downregulated genes in the HNRNPU_del/+_ -CTRL comparison; light blue: expressed in the dataset but not differentially expressed in the two conditions; light green: not expressed). In the figure HNRNPU_del/+_ samples are indicated as “HNRNPUdel”.

### Chromatin rewiring upon *HNRNPU* locus downregulation

Next, we performed Hi-C to investigate the changes in chromatin organization at a higher resolution (Lieberman-Aiden et al., 2009) . Since the NucleusJ analyses showed similar alterations of the heterochromatic fraction and nuclear shape in the two experimental conditions, we performed HiC on CTRL and HNRNPU_del/+_ cells at D0 and D28 (**Figure S6B-C and Methods**) as a model for HNRNPU haploinsufficiency. The number of chromatin interactions throughout the genome was similar at D0 between the two cell lines. However, at D28, a diverging pattern appeared in the chromatin organization maps, with the largest differences observed at the level of A/B compartments (10^5^–10^6^ base pairs) (**Figure S6D**). Furthermore, while the ratio of short and long-range contacts per chromosome was similar at D0, at D28 CTRL cells showed a higher ratio for the short- and long-range contacts than HNRNPU_del/+_ cells (**Figure S6E**). An analysis of differential chromosome contact frequencies between the HNRNPU_del/+_ and CTRL cells revealed little differences at D0 and higher contact differences at D28, as shown by the log2 ratio of the contacts at the chromosome level (**Figure 4B**).

Furthermore, a compartment interaction analysis revealed similar interaction strengths between the samples at D0 (compartmentalization strength: 1.30 and 1.28 for CTRL and HNRNPU_del/+_ cells, respectively). In contrast, we observed stronger compartmentalization in HNRNPU_del/+_ at D28 (1.31 vs. 1.59 for CTRL and HNRNPU_del/+_ cells, respectively), with higher interactions between active compartments (A compartment) and fewer trans-interactions compared to the CTRL cells (**Figure 4C**). These results align with the Hi-C experiments previously conducted on *Hnrnpu*-deficient mouse hepatocytes, showing decreased A-B interactions and increased A-A and B-B interactions upon *Hnrnpu* downregulation(Fan et al., 2018).

Next, we analyzed the compartment composition of each sample to identify genomic regions that switch compartments between CTRL and HNRNPU_del/+_ (**Methods**). At D0, 3% of the compartments switched from inactive (B) to active (A) compartment and 5.6% from A to B comparing CTRL and HNRNPU_del/+_, while at D28, 4.2% switched from B to A and 8.6% from A to B (**Figure 4D**). To investigate whether these compartment switches affect disorder-related genes, we assessed the enrichment for the genes mapping in genomic regions that switch compartments. Genes mapped in regions switching from A to B compartment were enriched in all the gene lists (ASD FDR: 0.0043; ID: 0.0018; Epilepsy: 0.0035, DD: 0.0136; hypergeometric test), whereas genes mapping in regions switching from B to A compartment were enriched only in the ASD gene list (FDR= 0.0136) (**Figure 4E and Supplementary Table 4**). In contrast, we found no significantly enriched GO terms.

Lastly, we investigated whether the compartment changes were related to the transcriptional changes described above. Therefore, we mapped the DEGs at D28 (|log2FC| > 0.58, Base Mean>20) with gene mapping in the switching compartments. The concordant genes were defined as genes mapping to a region with a compartment switching from A to B and downregulated in the transcriptome analysis, or genes mapping in the B to A compartment switch and upregulated. We identified 144 concordant genes at D0 and 241 concordant genes at D28 (**Supplementary Table 7**). Only concordant genes mapping in the compartment switching from A to B at D28 were significantly enriched for ASD (FDR: 0.014) and epilepsy (FDR: 0.007) gene lists (**Figure 4E and Supplementary Table 4**). By further analyzing the concordant genes to identify the enriched pathways, we showed that no ontologies were enriched in the concordant B to A genes, while the most significant enriched GO term in the concordant A to B genes was ‘GABA-gated chloride ion channel activity’ (FDR < 0.05) (**Figure S6F**), driven by genes such as *CACNB2*, *GABRA2*, *GABRA4*, *GABRB1*, *GABRG1,* and *SCN1A*. Interestingly, the *GABR* family genes downregulated in HNRNPU_del/+_ at D28 belong to a cluster on chromosome 4 with an approximately seven megabase (Mb) region that maps in the A compartment in CTRL and B compartment in HNRNPU_del/+_ at D28 (**Figure 4F**).

Overall, the Hi-C analysis showed that, similar to the transcriptome analyses, the compartment organization is more affected in HNRNPU_del/+_ at D28 than at D0, and *HNRNPU* deficiency led to an enrichment of B compartments at both timepoints. Interestingly, genes mapping in the enriched B compartments are associated with ASD and epilepsy, suggesting that the chromatin remodeling dependent on HNRNPU expression might ultimately be the driver of the observed phenotypes.

## DISCUSSION

Heterozygous genetic variants in the *HNRNPU* locus lead to various disorders with predominant brain phenotypes. Recently, few studies focused on the effects of mutations in *HNRNPU* gene in mouse and human cortical organoids. Here, we model microdeletions of the *HNRNPU* locus responsible of HNRNPU-related disorder and provide novel evidence of the molecular and cellular consequences of *HNRNPU* deficiency in human neuronal cells with hindbrain phenotype, adding knowledge on the effect *HNRNPU* deficiency on the brain region where *HNRNPU* expression is the highest.

We demonstrate that adequate levels of transcripts from the *HNRNPU* locus are needed in the early developmental transition from neural progenitors to developing neurons for proper neurogenesis. Our results, consistent with the earlier reports, show that *HNRNPU* expression is highest at early neural stem cell and progenitor stages and in a subpopulation of neural progenitors at the later stage of neuronal differentiation (Connolly et al., 2022; Sapir et al., 2022). Despite this high expression, results indicate that *HNRNPU* deficiency does not affect neural cells at this early stage.

In contrast, as the neural progenitor cells commit to differentiation, the reduced *HNRNPU* levels led to a higher number of dividing neural progenitors compared to the control stage, in a phase where the proportion of progenitor cells should decrease. This delayed transition from progenitors to differentiating neurons could explain the lack of differences in *HNRNPU* expression in HNRNPU_del/+_ from D0 and D5, and the almost equal expression levels with CTRL cells at D28, despite the heterozygous deletion of one allele. A recent study showed opposite effects for the complete loss of *Hnrnpu,* as it led to decreased proliferation followed by cell death of neural progenitors and postmitotic neurons in mice (Sapir et al., 2022). As heterozygous mutations in *HNRNPU* are not lethal, and the severity of the phenotypes in *HNRNPU*-related disorders is variable (Balasubramanian, 2022), embryonic cells can likely adapt to low levels of *HNRNPU* and still proliferate and differentiate. Therefore, it is reasonable that iPSCs and NES cells that retain 30–70% of the physiologic *HNRNPU* levels do not show a major phenotype, as we have shown here. Instead, we propose that cells with *HNRNPU* haploinsufficiency are inadequate to drive efficient cell fate transition of mitotic cells into maturing neurons and other neural cells through multiple regulatory pathways, leading to stochastic rewiring of the hindbrain development. Indeed, altered regulation of proliferation has been demonstrated to cause defects in the progenitors’ fate and, ultimately, neuronal development trajectories (Lalli et al., 2020; Pilaz et al., 2016). In several cellular models of ASD, unbalanced neural progenitors population due to both hyper- and hypo-proliferation of the progenitors have been documented (Connacher et al., 2022; Marchetto et al., 2017; Mowat et al., 2003; Zucco et al., 2018). Accordingly, we hypothesize that the observed downregulation of the synaptic and neuronal maturation markers is due to abnormal enrichment of progenitor at D28 stage and a consequence of the delayed maturation process. This observation is in contrast with what was observed in a recently published study on human cortical organoids, where *HNRNPU* mutations are shown to associate with downregulation of ontologies referring to nucleic acid binding and upregulation of neurogenic pathways (Ressler et al., 2023). The dysregulated genes are partially resembling transcriptomic alterations in embryonic mice carrying a heterozygous mutation in HNRNPU but are discordant with the perinatal mice. Thus, the stage of cell maturation and development in which the analyses are performed seem to be critical for studying effects of HNRNPU. Moreover, in this study, we are modeling the effect of the microdeletion of the whole *HNRNPU* locus on a hindbrain cell model, in contrast with single *HNRNPU* mutations on brain cortex systems, likely contributing to the discrepancy of the observed effects of HNRNPU mutations.

We provide mechanistic insights that both RNA processing and chromatin regulation in early brain development play a critical role in the observed brain phenotypes in *HNRNPU*-related disorders. It has been earlier demonstrated that the correct pool of mRNA isoforms from the alternative splicing process is important in the transition from progenitor cells to neurons in the developing cerebral cortex (Zhang et al., 2016), and RNA splicing is one of the key enriched pathways from molecular and genetic studies of ASD (Gandal et al., 2018; Satterstrom et al., 2020). Therefore, our and others’ results pinpoint that further delineation of the RNA splicing program during the early steps of brain development is essential for understanding the origins of NDDs.

Furthermore, the importance of 3D genome organization for cell fate decisions during neural development is starting to emerge, showing that dynamic changes at multiple levels of chromatin organization are needed for these processes (Bonev et al., 2017; Hu et al., 2021). This is in line with our results showing major reorganizations in a later stage of neural differentiation. Indeed, multiple NDD cell models have shown that the dysfunction of the chromatin organization leads to changed neuronal maturation (Calzari et al., 2020; Markenscoff-Papadimitriou et al., 2021; Parisian et al., 2020; Su et al., 2021).

We additionally provide evidence of cellular processes, such as cilia organization and synaptogenesis, that are affected by the molecular changes in the *HNRNPU* deficiency state. For instance, we demonstrate an increase in ciliated cells. Recently, HNRNPU was indicated to localize occasionally to cilium in mice brain cells (Sapir et al., 2022). As *HNRNPU*-related disorders share many phenotypic features with ciliopathies (Balasubramanian, 2022; Focșa et al., 2021), the connection between cilia organization and *HNRNPU* warrants more studies.

In conclusion, we provide the first evidence of the functional consequences of *HNRNPU* mutations in human neural progenitors in the hindbrain in the form of an inadequate switch to neurogenesis. This, in turn, leads to large-scale effects on chromatin organization and transcriptional landscape at later stages of neural development and presumably to diverging trajectories of neurons and other neural cells. Follow-up studies of direct targets, different developmental stages, and brain regions using both 2D and organoid models will be needed to assess better the impact of *HNRNPU* haploinsufficiency on neurogenesis and its role in the pathogenesis of *HNRNPU*-related disorders.

## LIMITATIONS OF THE STUDY

We acknowledge that this study has limitations, including that we only compare the isogenic model constructed with siRNA with one patient cell line, providing the limited possibility to analyze the genetic background effects, recently shown to be highly important to study(Paulsen et al., 2020). Moreover, while mimicking the heterozygous deletion of the whole locus, we cannot distinguish if the observed effects are mainly driven by HNRNPU or if they are the results of a combined role of HNRNPU and *HNRNPU-AS1*. Furthermore, we focused on a very early model of neural development using NES cells and undirected differentiation, which cannot represent the complexity of the human hindbrain. The analyses described here are from a pool of cells not sorted for the cell type or analyzed as single cells, thus, we only give a general overview of the transcriptional and chromatin organization landscape related to HNRNPU haploinsufficiency. Further studies using single-cell techniques, different neuronal differentiation models and hindbrain human organoids should be employed to present the cell-specific effects of HNRNPU mutations in later stages of brain development.

## ACKNOWLEDGMENTS

The authors are grateful to the donors of the cell lines for their participation in this research. The authors acknowledge support from the National Genomics Infrastructure in Stockholm funded by Science for Life Laboratory, the Knut and Alice Wallenberg Foundation and the Swedish Research Council. The computations and data handling were enabled by resources provided by the National Academic Infrastructure for Supercomputing in Sweden (NAISS) and the Swedish National Infrastructure for Computing (SNIC) at Uppsala Multidisciplinary Center for Advanced Computational Science, partially funded by the Swedish Research Council through grant agreements no. 2022-06725 and no. 2018-05973. We thank A. Smialowska at NBIS (National Bioinformatics Infrastructure Sweden) for technical support with HiC data analysis, which was made possible through application support provided by SNIC. The authors acknowledge support from the iPS Core facility at Karolinska Institutet for assistance with the generation of iPSC and NES cells. The project was supported by grants from the Swedish Research Council to K.T. (grant no. 2017-01660), S.B. (grant no. 2019-01303), and N.C. (grant no. 2018-02950); Swedish Foundation for Strategic Research to K.T. (grant no. FFL18-0104); the Swedish Brain Foundation – Hjärnfonden to K.T.; the Karolinska Institutet KID Program to N.C., KI-NIH PhD program for PhD students (M.O.), Osk. Huttunen Foundation (M.O.), Åke Wiberg Foundation to K.T.; the H.K.H. Kronprinsessan Lovisas förening för barnasjukvård och Stiftelsen Axel Tielmans minnesfond to F.M.; the Swedish Foundation for Medical Research (SSMF) to F.A.; The Foundations of Petrus och Augusta Hedlund to K.T.; the Strategic Research Area Neuroscience (StratNeuro) at Karolinska Institutet to K.T.; the Swedish Foundation for International Cooperation in Research and Higher Education (STINT) to K.T.; and the Committee for Research at Karolinska Institutet to K.T., grant #R35GM128661 from the National Institutes of Health to Y.J. (grant #R35GM128661). Open access funding is provided by Karolinska Institutet.

## AUTHOR CONTRIBUTIONS

Conceptualization: F.M., M.O., and K.T.; Methodology and Data Analysis: F.M., M.O., C.S., S.A., A.A., R.B., M.M., F.A., A.P., M.W., I.R., M.B., D.L., B-M.A., J.I., K.L.R., Mo.M., Y.J., A.F., A.B., and K.T.; Validation: F.M., M.O., C.S., S.A., A.A., and K.T.; Resources: S.B. and K.T.; Data Curation: F.M., M.O., A.A., F.A., A.P., M.W., D.L., B.M.A., J.I., K.L.R.; Writing – Original Draft: F.M. and K.T.; Writing – Review & Editing: F.M., M.O., and K.T.; Visualization: F.M., M.O., D.L., and K.T.; Supervision: Y.J., N.C., M.Bi., E.S., A.B., and K.T.; Project Administration: F.M. and K.T.; Funding Acquisition: F.M., Y.J., N.C., E.S., A.B., S.B., and K.T., Critical review of the draft: All authors.

## SUPPLEMENTARY FIGURES

**Figure S1.**
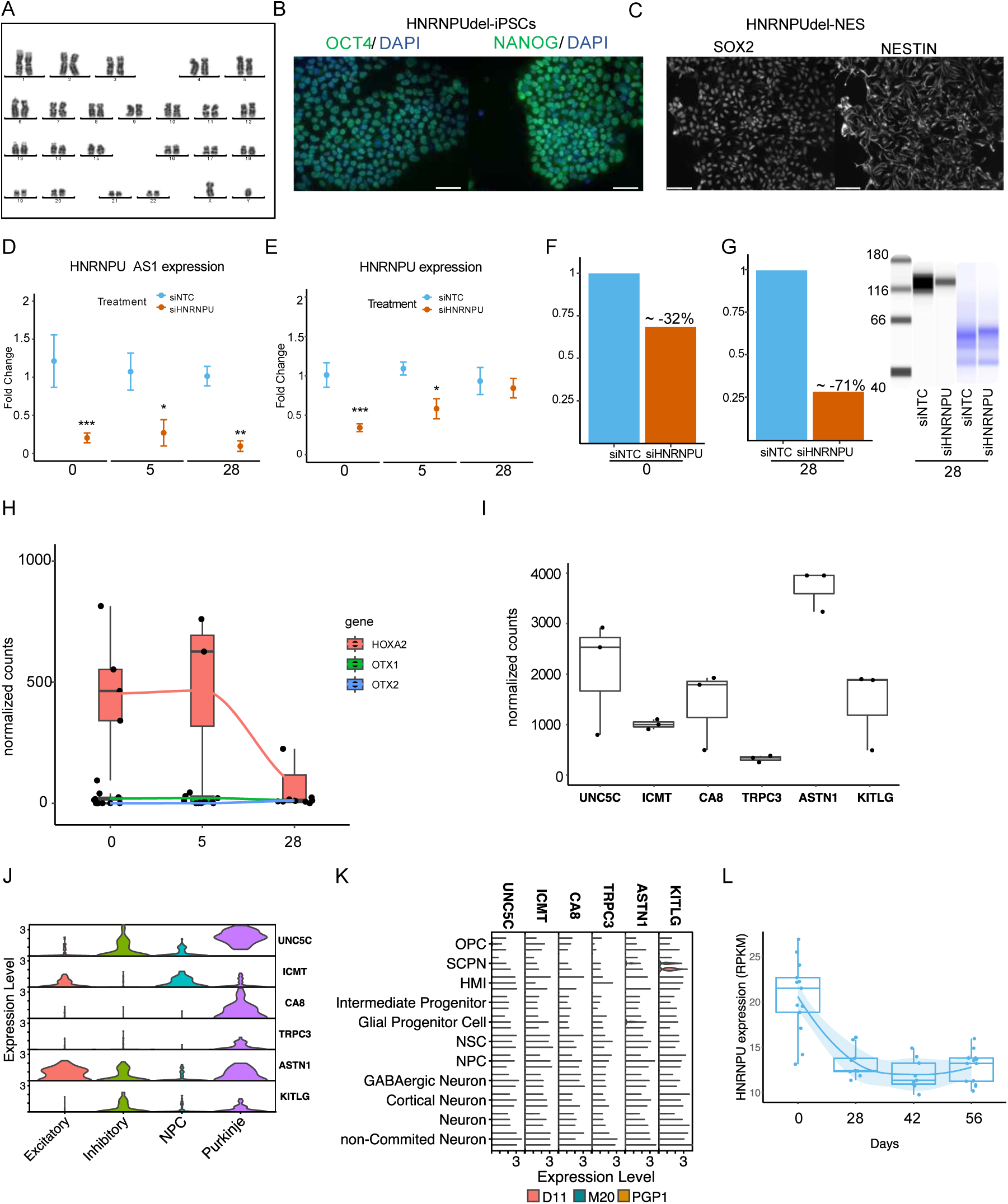
Characterization of the two *HNRNPU* knockdown conditions. **A)** Karyotype of iPSCs obtained from the fibroblasts of the individual carrying the heterozygous deletion of HNRNPU, and from which the HNRNPU_del/+_ cells were derived. **B)** Staining the HNRNPU_del/+_ - iPSC cells with the pluripotency markers OCT4 and NANOG. Nuclei are counterstained with DAPI. Scale bar= 50µm. **C)** Immunostaining of SOX2 and NESTIN in HNRNPU_del/+_ cells at D0. Scale bar= 50µm. **D-E)** *HNRNPU-AS1* (**D**) and *HNRNPU* (**E**) RNA expression after treatment with siHNRNPU at D0, D5 and D28. Bar plot normalized for each siNTC sample. **F-G)** HNRNPU protein expression after treatment with siHNRNPU at D0 (**F**) and D28 (**G**). Bar plot normalized for each siNTC sample. Capillary western blot representation (right) of the samples at D28 probed with antibody against HNRNPU and total protein quantification in the following lanes. **H)** Expression of embryonic markers *HOXA2*, *OTX1*, *OTX2* in CTRL cells at D0, D5 and D28. **I)** Expression of cerebellar markers (*UNC5C*, *ICMT*, *CA8*, *TRPC3*, *ASTN1*, *KITLG)* in CTRL cell line at D28. **J)** Expression of cerebellar markers among subtypes of neuronal cells in our previously published scRNA-seq data(Becker et al., 2020) at D28 visualized in violin plot. NPC=Neural progenitor cell. **K)** Expression of cerebellar markers among cell types and across samples in organoid scRNA-seq data(Ressler et al., 2023). D11=*HNRNPU* deficient organoids with a heterozygous frameshift mutation, M20=*HNRNPU* deficient organoids with a heterozygous premature termination codon, PGP1=Control organoids. **L)** *HNRNPU* RNA expression in neurotypical samples from published iPSC-neuronal study dataset(Burke et al., 2020). The day 0 refers to rosette forming cells comparable to NES cells and the following time points are days in differentiation.

**Figure S2.**
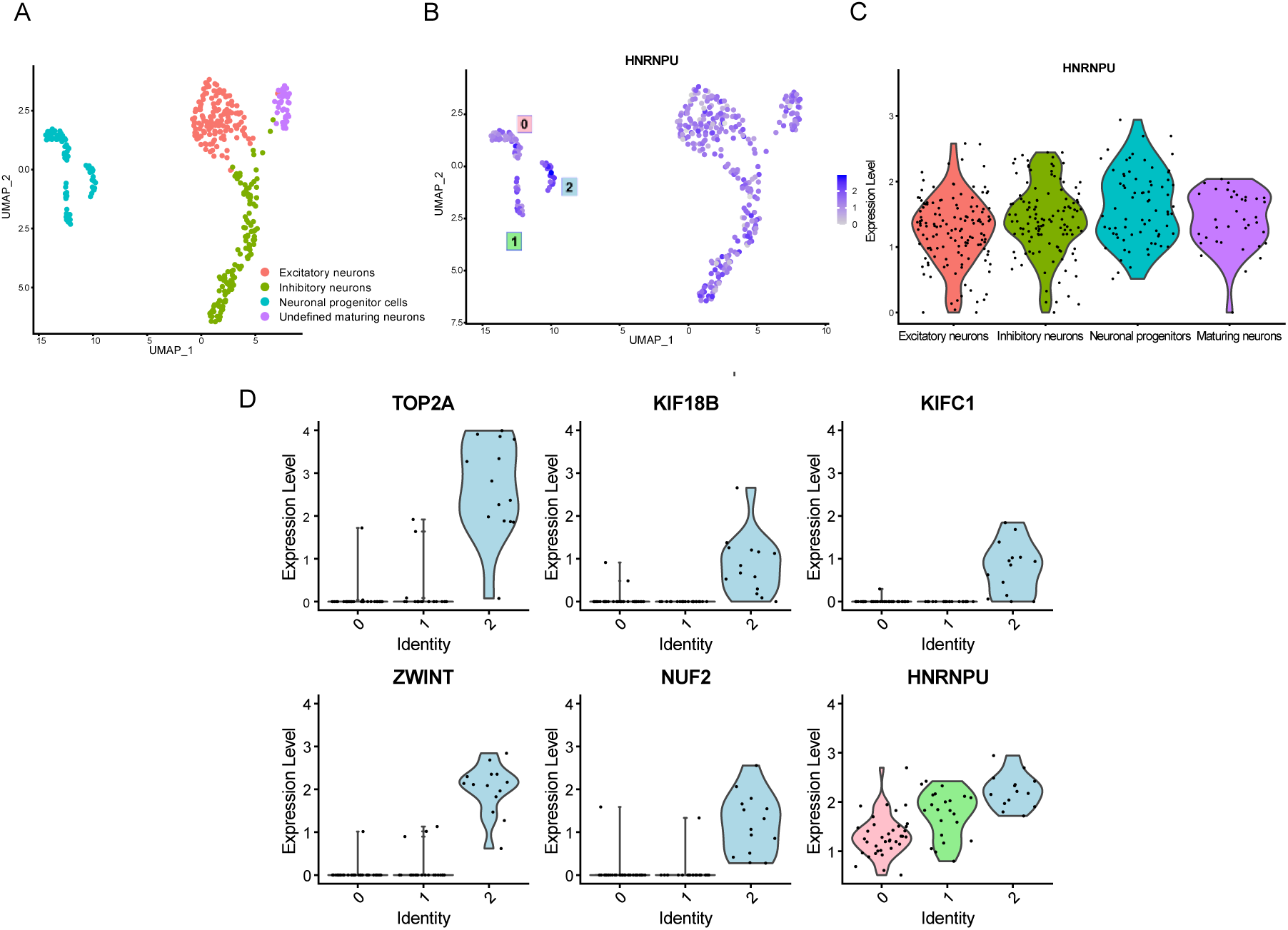
*HNRNPU* expression is higher in proliferating progenitor cells. **A)** UMAP dimensional reduction of scRNA-seq data from previously published cells at D28, obtained with the same differentiation protocol as the cell model in study(Becker et al., 2020). The different cell type populations are highlighted. **B)** Feature plot of HNRNPU expression in the cell populations from the scRNA-seq analysis **C)** Violin plot of HNRNPU expression in each cell type from the scRNA-seq analysis. **D)** Expression of proliferation markers in the subclusters defining the NPCs subpopulation from the scRNA-seq analysis. 0-1-2 refer to the NPCs subpopulations showed in H. In the figure HNRNPU_del/+_ samples are indicated as “HNRNPUdel”.

**Figure S3.**
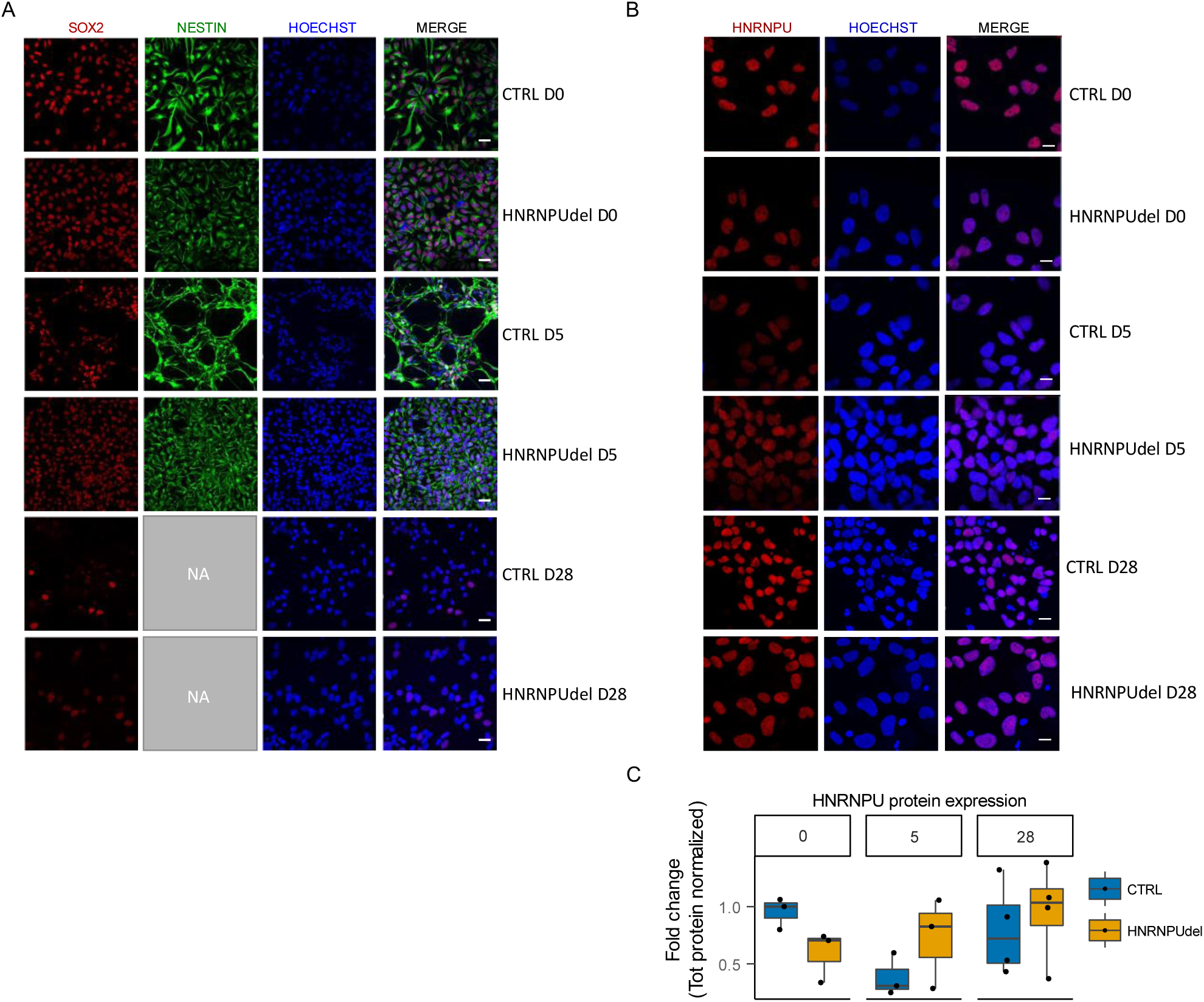
HNRNPU protein localization and expression during neuronal differentiation. **A)** Immunostaining of HNRNPU in red, Hoechst in blue and merge at D0, D5 and D28 in CTRL and HNRNPU_del/+_. Scale bar= 10 µm. **B)** HNRNPU protein quantification during differentiation in CTRL and HNRNPU_del/+_. In the figure HNRNPU_del/+_ samples are indicated as “HNRNPUdel”.

**Figure S3.**
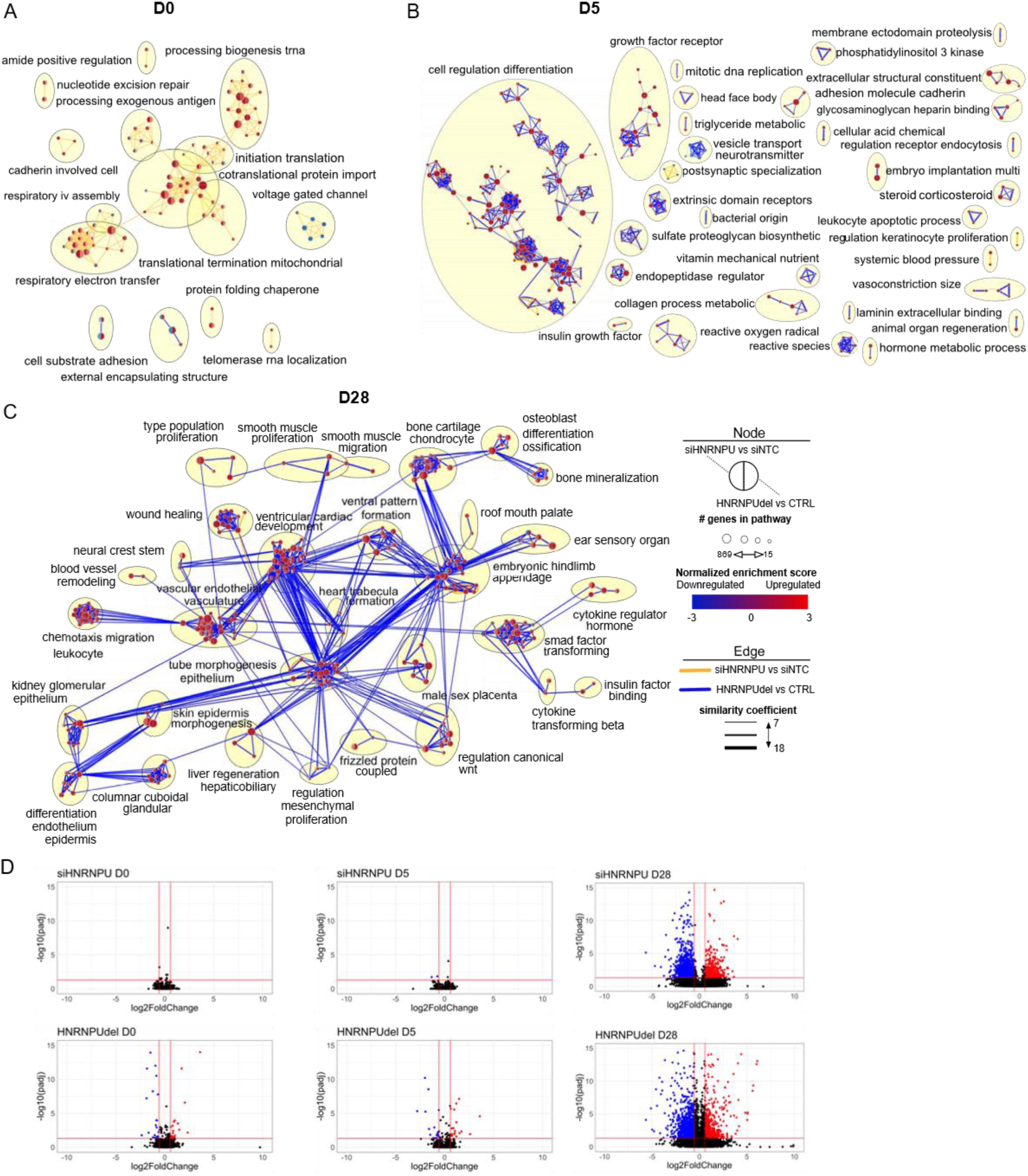
Extended differential pathway enrichment under HNRNPU downregulation during neurodevelopment. **A-B)** All the upregulated and downregulated pathways, respectively, at D0 and D5. **C)** Upregulated pathways at D28 related to general development. For each node, the left half indicates the siHNRNPU versus siNTC enrichment and the right half the HNRNPU_del/+_ versus CTRL enrichment. The color of the edge indicates which of the datasets significantly contributed to the pathway call. **D)** Volcano plots of DEU exon bins at D0, D5 and D28 for both HNRNPU deficient conditions. The plots are zoomed in, and some highly significant exon bins are thus excluded from the D28 plots to enhance readability. In red is the upregulated DEU, and in blue is the downregulated DEU. The vertical lines indicate the threshold for significance (p adj < 0.05 and |log2FoldChange|>0.58). In the figure HNRNPU_del/+_ samples are indicated as “HNRNPUdel”.

**Figure S4.**
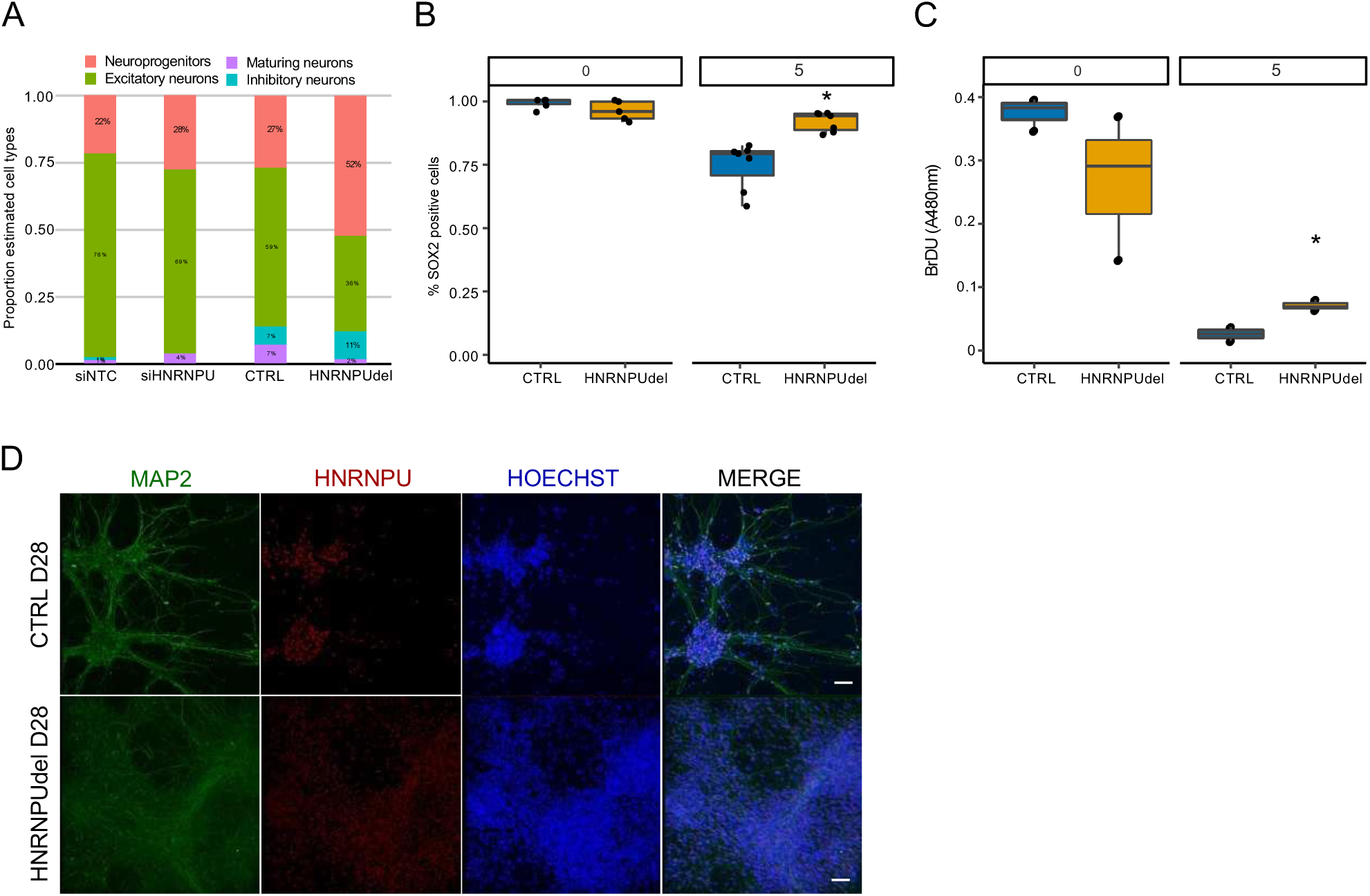
Delayed maturation and higher progenitor phenotype under HNRNPU downregulation during the differentiation time course. **A)** Estimated proportion of the different cell types at D28 after deconvolution analysis. **B)** SOX2 positive cells at D0 and D5 in CTRL and HNRNPU_del/+_. **C)** BrdU signal at D0 and D5 in CTRL and HNRNPU_del/+_. **D)** MAP2, HNRNPU and Hoechst staining of CTRL and HNRNPU_del/+_ at D28. Scale bar= 50µm. *p<0.05. In the figure HNRNPU_del/+_ samples are indicated as “HNRNPUdel”.

**Figure S5.**
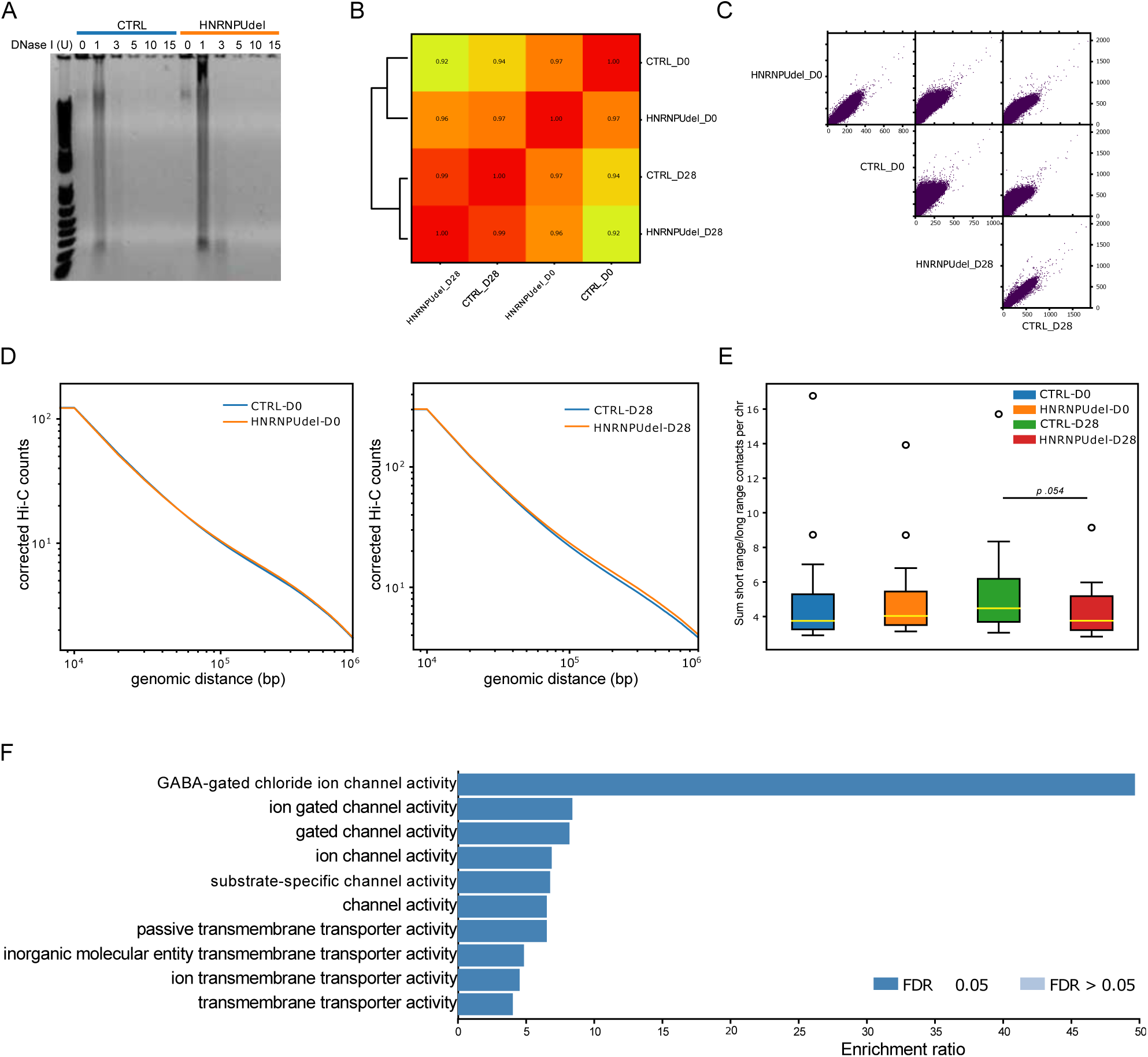
Extended effects of HNRNPU downregulation on chromatin organization. **A)** DNaseI sensitivity assay in CTRL and HNRNPU_del/+._ **B)** Heatmap of the correlation matrixes of the HiC samples. **C)** Pearson’s correlation matrixes of the HiC samples. **D)** Corrected HiC counts at a specific genomic distance at D0 (left) and D28 (right). **E)** Boxplot of short range/long range HiC contact per chromosome. **F)** Pathways enriched in the “concordant A to B” genes, according to Over-representation analysis from WebGestalt. In the figure HNRNPU_del/+_ samples are indicated as “HNRNPUdel”.

## KEY RESOURCES TABLE

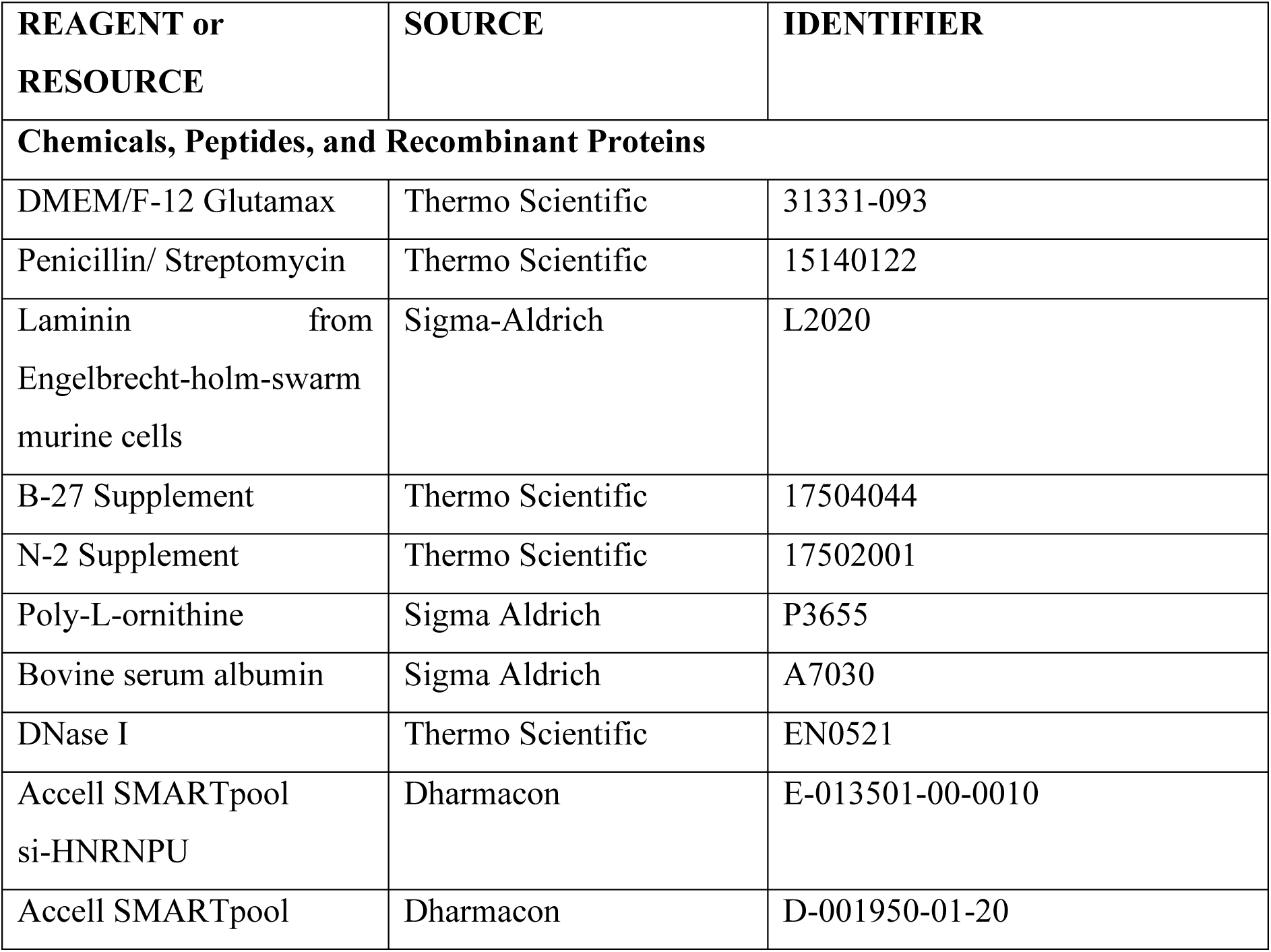
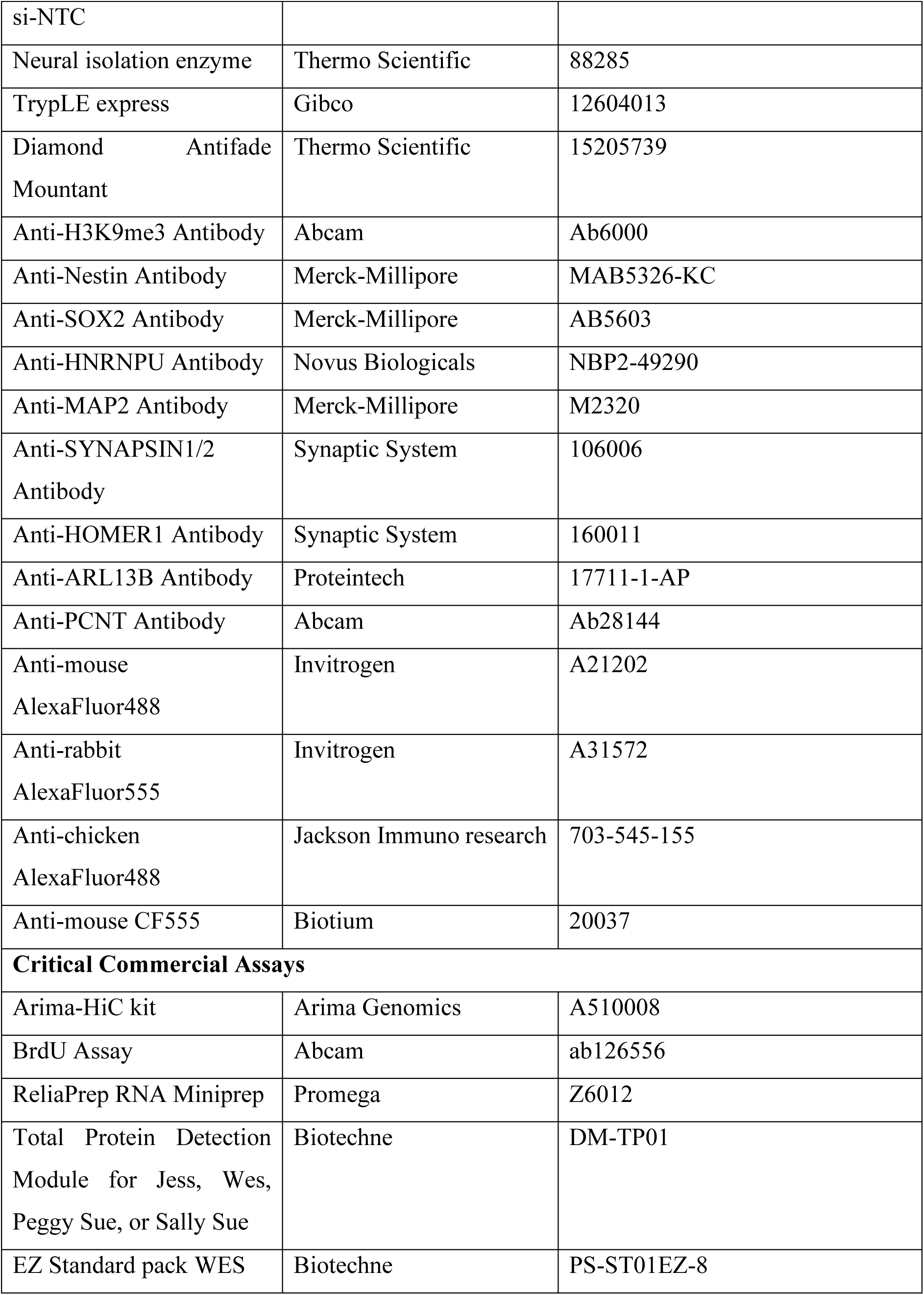
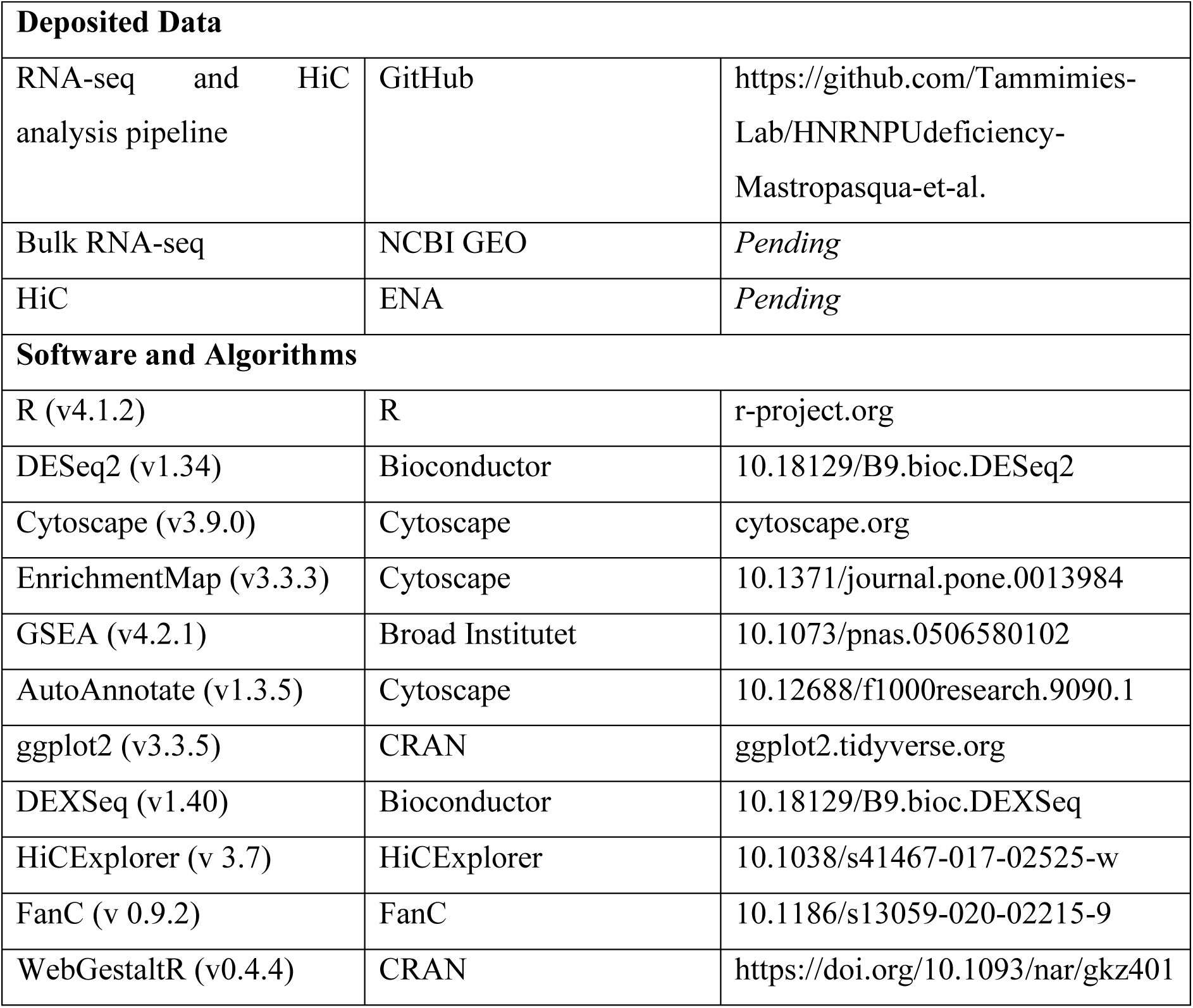

## LEAD CONTACT AND MATERIALS AVAILABILITY

Further information and requests for resources and reagents should be directed to and fulfilled by the Lead Contact, Kristiina Tammimies.

## DATA AND CODE AVAILABILITY

All the related code and data matrixes are available in the GitHub https://github.com/Tammimies-Lab/HNRNPUdeficiency-Mastropasqua-et-al RNA-seq and Hi-C data will be uploaded to GEO and ENA.

## METHODS

### Identification of the individuals with *HNRNPU* deletion and phenotypic characterization

We have earlier performed a screening for copy number variation in a twin sample from The Roots of Autism and ADHD study in Sweden (RATSS) (Bölte et al., 2014; Stamouli et al., 2018) in which we identified a male twin pair carrying the deletion on chr1:244997953-245042312 (hg19) encompassing *HNRNPU*. The twin pair has undergone an extensive phenotypic characterization, including evaluation for ASD, ID, and other NDDs, cognitive testing, and medical examination (Supplementary table 1). Written informed consent was obtained from individuals and their parents prior to the study. The study was approved by the regional and national ethical boards in Sweden and has been conducted in accordance with the Declaration of Helsinki for medical research involving human subjects, including research on identifiable human material and data.

### Cell culture

Human iPSC cells were derived from fibroblasts of a male carrying a 44Kb mutation of the HNRNPU locus spanning through *COX20*, *AS1-HNRNPU,* and *HNRNPU*, using a previously described protocol (Uhlin et al., 2017b). Human iPSC cells were cultured in mTeSR Plus (Stem Cell Technology) on 5 µg/ml BioLaminin 521LN (Biolamina) precoated vessels. Pluripotency and normal karyotype were confirmed, as shown in Supplementary Fig 1. Dual-SMAD inhibition was applied to derive neuroepithelial stem (NES) cells from human iPS cells as described previously (Chambers et al., 2009; Falk et al., 2012). Patient-derived and previously established NES cells from a male neurotypical donor (Uhlin et al., 2017a) were seeded on 20 µg/ml poly-L-ornithine (Sigma Aldrich), and 1 µg/ml laminin (Sigma Aldrich) precoated plastic surfaces in DMEM/F12 + Glutamax medium (Gibco) supplemented with 0.05X B-27 (Gibco), 1X N-2 (Gibco), 10 ng/ml bFGF (Life Technologies), 10 ng/ml EGF (PeproTech) and 10 U/ml penicillin/streptomycin (Gibco), replacing media every day. For immunofluorescence, the glass coverslips were precoated with increased concentrations of poly-L-ornithine to 100 µg/ml and laminin to 2 µg/ml. Cells were maintained in a 5% CO_2_ atmosphere at 37°C. Upon starting the differentiation, cells were changed to medium with an increased 0.5X B-27 concentration without bFGF and EGF. Two-thirds of media supplemented with 0.4 ug/ml laminin was changed every other day until differentiation 15; whereafter media was changed every third day. NES cells were harvested two days after culturing and neural cells were harvested after 5, 28 or 50 days of differentiation.

### DNAse I sensitivity assay

Cells were grown and harvested as earlier described and resuspended in cold RSB buffer: 10mM Tris-HCl (pH7,4), 10mM NaCl, and 3mM MgCl_2._ Cells were then lysed in cold lysis buffer (RSB buffer and 0,2% Triton X-100) and centrifuged to collect nuclei in the pellet. The nuclei were then incubated with different amounts of DNase I enzyme (0, 1, 3, 5, 0, 15 units) at 37°C for 10 min. The digestion was blocked by adding 50mM of EDTA followed by incubation at 55◦C for 1min. The results of the digestion were visualized on an 0.8% agarose gel.

### siRNA-mediated silencing

Cells were transfected with 0.5 µM Accell SMARTpool siRNA targeting *HNRNPU* mRNA (Dharmacon E-013501-00-0010) or 0.5 µM Accell Nontargeting siRNA (Dharmacon D-001950-01-20) according to manufacturer’s protocol. The siRNA pool consisted of the following oligos: oligo1: UCUUGAUACUUAUAAUUGU, oligo2: CUCGUAUGCUAAGAAUGGA, oligo3: GUUUCAGGUUUUGAUGCUA, oligo4: CUAGUGUGCUUGUAGUAGU. NES cells were transfected one day after seeding, and samples were collected 24 hours after treatment. Cells under differentiation were transfected once in the NES phase, once when changing media to differentiation media, and thereafter every sixth day until D28 sample harvesting.

### Immunofluorescence and image analysis

Cells on glass coverslips were fixed in 4% formaldehyde for 20 minutes at room temperature and washed with 1X TBS. Blocking was performed with 5% Donkey Serum and 0.1% Triton X-100 in 1X TBS for 1 hour. Primary antibodies diluted in blocking buffer (Nestin 1:1000; SOX2 1:1000; Synapsin ½ 1:500; Homer1 1:250; MAP2 1:500; HNRNPU 1:500; H3K9me3 1:500, ARL13B 1:10000, PCNT 1:250) were incubated at +4°C overnight. Careful washing was done with 1X TBS, and secondary antibodies diluted 1:1000 in blocking buffer were incubated at room temperature for one hour, covered from light. Coverslips were washed and mounted with Diamond Antifade Mountant (Thermo Scientific). Images were acquired with LSM 700 Zeiss Confocal Microscope using 20x or 63x magnification and 0.5 µm z-stack through the sample.

At least three replicates per sample were stained and analyzed for the immunocytochemistry analyses. The images stained with synapsin 1/2 were analyzed with ImageJ plugin Synapse Counter, and the statistical Student T-test analyses were performed in R. The statistical analyses for SOX2 positive nuclei were performed in R by ꭓ^2^-test. For primary cilia analysis, a cell was considered ciliated if positive for both cilia markers ARL13B and PCNT; the statistical testing was performed in R by ꭓ^2^-test. For the images of the nuclei stained with the H3K9me3 antibody, the single nuclei were cropped using an in-house built macro for ImageJ. A minimum of thirty nuclei per replicate was analyzed with NucleusJ, a plugin of ImageJ, using the default settings (Poulet et al., 2015).

### Capillary Western blot

Cells were collected in extraction buffer (50 mM Tris-HCl, 100 mM NaCl, 5 mM EDTA, 1 mM EGTA) supplemented with a 1X protease inhibitor cocktail (Thermo Scientific) using a plastic cell scraper. The samples were sonicated with six short pulses at 36% amplitude (Vibra-Cell VCX-600, Sonics). Protein concentrations were measured with Qubit Protein Assay Kit (Thermo Scientific). The total protein sample (150 ng/µl) was loaded on the capillary western blot system Simple-Western-JESS (Bio Techne), multiplexed for total protein and chemiluminescence detection of HNRNPU (1:10 dilution). Data were analyzed using Compass for S.W. software (5.0.1), and the HNRNPU peak area was normalized against the total protein area. Three to five biological replicates were analyzed for each time point, and the significance at each time point was evaluated by Student T-test.

### RNA extraction and bulk RNA sequencing analyses

Cells were lysed in TRIzol reagent (Invitrogen), and RNA isolated with ReliaPrep RNA Cell Miniprep kit (Promega Z6012). We extracted RNA samples for three to five biological replicates per cell line and time point. Samples were delivered to NGI Sweden for library preparation and sequencing. The samples were subjected to library preparation with Illumina Truseq Stranded total RNA RiboZero GOLD kit, except for siNTC-D28 and siHNRNPU-D28 libraries prepared with Illumina TruSeq Stranded mRNA kit due to low RNA yields. All libraries were sequenced on the NovaSeq6000 platform with a 2×151 setup using NovaSeqXp workflow in S4 mode flowcell. We obtained, on average, 35 and 65 million reads per sample for early time points and D28, respectively, with a minimum 82,7% alignment rate.

Differential gene expression analysis was performed using DESeq2 (v1.24.0) (Love et al., 2014) in R (v4.1.2). The significance thresholds used were adjusted *p* value < 0.05 (Benjamini-Hochberg adjustment), base mean > 20, and absolute log2 fold change > 0.58. Pathway analysis was done according to previously described protocols (Reimand et al., 2019). In short, the ranked gene expression list was used in Gene set enrichment analysis (GSEA) (Version 4.3.0) to analyze the enrichment in the gene ontology, molecular function, and biological process gene lists (v. 7.4), and enriched categories were visualized in Cytoscape (v3.8.2) with Enrichment Map (v3.1.0) and AutoAnnotate (Merico et al., 2010; Reimand et al., 2019; Shannon et al., 2003).

Differential exon usage analysis was performed using the DEXSeq package (v1.40.0) (Anders et al., 2012). A flattened annotation file was created using provided python script, excluding the aggregate exon bins, and exon counts were counted using provided python script. The analysis was performed with the formula *∼ sample + exon + condition: exon.* A gene was called to have evidence for DEU if at least one exon bin was differentially used between conditions. The difference was considered significant with FDR (Benjamini-Hochberg adjustment) adjusted P<0.05, exon base mean >10, and an absolute log2 fold change>0.58. Over-representation analysis (ORA) was performed using the online tool WebGestalt (Liao et al., 2019). In addition, hypergeometric tests were used to test for enrichment between differentially expressed genes and DEU genes and specific NDD-related gene lists: ASD gene list (SFARI genes selected for score 1,2 and syndromic, release 07-20-2022), ID gene list (green and amber genes from https://panelapp.genomicsengland.co.uk/ v3.1632), epilepsy gene list (green and amber genes from https://panelapp.genomicsengland.co.uk/ v2.547) and general development (compiled gene list (Becker et al., 2020)).

### Real Time PCR

The RNA was reverse-transcribed using iScript cDNA Synthesis Kit (BioRad) and cDNA quantified with SsoAdvanced Universal SYBR Green Supermix (BioRad) following the manufacturer’s protocols on a CFX96 thermal cycler (BioRad). The primer used are:

HNRNPU-AS1 (AGGAAGCTGTACACTGGAGG, CAATGTCTTCACCAATAACAAAGC); HNRNPU (AGTTTAACAGAGGTGGTGGCC, GCCCCTCCTATTATATCCGCC); GAPDH (AAGGTGAAGGTCGGAGTCAAC, GGGGTCATTGATGGCAACAATA). CFX Manager software was used to record amplification curves and to determine Ct values. RT-qPCR reactions were performed in technical triplicates. We calculated the ΔCt to the GAPDH housekeeping gene and ΔΔCt to control cell lines. We used three biological replicates of cells seeded at different passages. Statistical significance between cell lines was determined with ANOVA and post hoc Tukey HSD in R (v. 4.1.2).

### Hi-C sequencing

Cells were cultured as NES for D0 collection or differentiated for 28 days and harvested by briefly rinsing the cells in accutase and then incubating them with TrypLE express (Gibco) and neural isolation enzyme (Thermo Scientific). Samples were then processed using the Arima-HiC kit (PMID: 29779944) according to the Arima Genomics User Guide for Mammalian Cell Lines (Catalog number A510008). Briefly, we crosslinked harvested cells with 2% formaldehyde and then used approximately 1 million fixed cells as input for each replicate sample. Subsequently, we used 1.5 μg of Hi-C template for biotin pull-down and library preparation according to the Arima Genomics User Guide for Library Preparation using KAPA Hyper Prep Kit. Specifically, we used 8 PCR cycles for library amplification. We then sequenced the Arima-HiC libraries for each time point on one flowcell on the Illumina NovaSeq S Prime system, obtaining an average of 900M sequencing reads for D0 samples, and 1000M sequencing reads per sample for D28 samples. Sequencing was carried out at the National Genomics Infrastructure at the Science for Life Laboratory (SciLifeLab) in Stockholm, Sweden.

We processed the raw sequencing reads using the HiCUP pipeline (v0.7.4) (PMID: 26835000) with default parameters. Briefly, the pipeline employs Bowtie2 (v2.4.1) (PMID: 22388286) to align the reads to the human reference genome (GRCh37/hg19) and filter out experimental artifacts (i.e., circularised, re-ligated, and duplicate reads). We generated the ‘digest file’ using the ‘hicup_digester’ command with the Arima option (--Arima). We used the HiCUP output files (BAM format), which contain only valid, non-redundant read pairs, as input for pairtools (v0.3.0). First, we converted the BAM files into .pairsam format using the pairtools parse and sort modules. In addition to individual replicates, we generated pooled samples for each developmental stage by merging replicates using the pairtools merge module. We marked read duplicates using the pairtools dedup module, with option --mark-dups, and filtered the results by selecting only specific pair types (i.e., ‘U.U.’, ‘U.R.’ and ‘R.U.’) via the pairtools select module, which produced a .pairs format output. Finally, we added the fragment information using the ‘fragment_4dnpairs.pl’ convenience script provided alongside the Juicer pipeline and converted the .pairs files into .hic format using the Pre module from Juicer-Tools (v1.22.01). Unless explicitly required by the software/package, the contact matrices were normalized to the smallest library and corrected with ICE using HiCexplorer (v 3.7) (Ramírez et al., 2017). HiCexplorer package was used for mapping genomic contacts and contact enrichment between the samples.

The compartments were called at 1 Mb resolution, and the first four PCAs were extracted by HiCexplorer using the hicPCA tool and the gene track from hg19 to assure the correct orientation of the eigenvector. The PCA for each chromosome and each sample were chosen by considering the highest correlating PCA between the samples and the best mapping with the gene density eigenvector. The compartments were defined as A and B according to the eigenvector orientation, with A being the gene-rich and positive eigenvector and B being the gene-poor regions and negative eigenvector. FanC toolkit (v. 0.9.1) (Kruse et al., 2020) was used to calculate compartment strength and cis-trans-compartment interactions. The R package annotatr was used to annotate the genomic regions switching compartments in the different conditions.

### Deconvolution

The deconvolution of bulk RNA-seq data to previously published scRNA-seq sample (Becker et al., 2020) was done using the Bseq-SC package (Baron et al., 2016). Bseq-SC uses cell-type-specific marker genes from single-cell RNA transcriptomes to predict cell-type proportions underlying bulk RNA transcriptomes. Deconvolution was done with 30 marker genes for each cell population, except for cluster “Undefined maturing neurons”, which had only 25 defining genes. Statistical differences in the estimated cell proportions were calculated by the ꭓ^2^ statistical test in R (v4.1.2).

### Public data processing

Data from Human Brain Transcriptome (https://hbatlas.org) were obtained to visualize *HNRNPU* expression in different human brain regions during development. Additional stem cell time-course data was accessed via LIBD Stem Cell Browser (http://stemcell.libd.org/scb). RPKM values for HNRNPU were extracted for evaluating *HNRNPU* expression during neuronal differentiation from iPSCs (Burke et al., 2020). Seurat object for human cortical organoid data was downloaded from GEO with accession number GSE219317(Ressler et al., 2023).

### Cell proliferation assay

To analyze differences in cellular proliferation, the BrdU assay (ab126556, Abcam) was performed for three replicates at D0, D5, and D28 time points. Twenty-four hours prior to the read-out, 1X-BrdU reagent was added to the cell culture vessels for incorporation by incubation at 37°C and 5% CO_2_. The culture media was aspirated for the read-out, and cells were fixed with the supplied fixing solution. This was followed by exposure to the anti-BrdU antibody (primary antibody), incubation at room temperature for 1 hour, and washing with the supplied plate wash buffer. The cells were incubated with the HRP-tagged secondary antibody at room temperature for 30 minutes, followed by TMB exposure and recording absorbance at 450 nm. Afterwards, the cells were incubated in TBS, 0.1% triton-X 100, and Hoechst 1:1000 for 15 min, followed by fluorescence measurement, which was used to calibrate the BrdU quantification for the number of nuclei in each sample. The quantification of proliferation rate followed by Student T-test statistical comparison was done in R (v4.1.2).

### Electrophysiology

Whole-cell patch-clamp recordings were performed at 5, 8, 28, and 50 days of differentiation. Immediately prior to recordings, the cells were washed with 1X PBS, and then Krebs-Ringer’s solution composed of (in mM): NaCl 119, KCl 2.5, NaH2PO4 1, CaCl2·2H2O 2.5, MgCl2·6H2O 1.3, HEPES 20, D-Glucose 11 (pH 7.4) + Laminin 1:1000 was added in the dish. The recordings were performed in Krebs-Ringer’s solution. Recording pipettes were fabricated with a Narishige pc-100 puller and had resistances of 3-5 MOhm when filled with the internal solution composed of (in mM): 120 K-gluconate, 0.1 EGTA, 4 MgATP, 0.3 Na2GTP, 10 HEPES, 20 KCl and 5 Na2-phosphocreatine (pH 7.4). Current and voltage responses were measured at room temperature using a MultiClamp 700B amplifier (Molecular Devices) and digitized with Axon™ Digidata® 1550B analog-to-digital converter connected to a personal computer running pClamp 11.0.3 (Molecular Devices). Membrane capacitance and resistance were derived from the pClamp 11.0.3 (Molecular Devices) membrane-test function. Data analysis was performed by Clampfit 10.7 (Molecular Devices).

## SUPPLEMENTAL INFORMATION

Supplemental information can be found online.

