## Supplemental Table 1 for "Deficiency of the Heterogeneous Nuclear Ribonucleoprotein U locus leads to delayed hindbrain neurogenesis"

**Supplementary table 1**. Phenotype information of monozygotic twin pair carrying heterozygous deletion affecting HNRNPU locus

| | Twin 1 | Twin 2 |
|---|---|---|
| Birth week and delivery mode | 36 + 4 with caesarean section | |
| Birth measures | Weight: 2310 g, height: 44 cm<br>Head circumference: 34<br>Apgar score: 9, 9, 10 | Weight: 2340g, height: 44 cm<br>Head circumference: 33<br>Apgar score: 8, 10, 10 |
| Problems in infancy | Feeding problems<br>Jaundice<br>Frequent ear infections<br>Eczema<br>Scarlet fever<br>Phototherapy after birth | Feeding problems<br>Jaundice<br>Frequent ear infections<br>Eczema<br>Scarlet fever<br>Oxygen therapy after birth |
| Early development | Language: Delayed (first words 84 months old, sentences 116 months)<br>Motoric: Delayed (walked at 4 year of age)<br>Food: Problems with breast-feeding and with solid food<br>Toilet training delayed (not obtained by 9 years of age)<br>First noted abnormalities: 6 months | Language: Delayed (first words 54 months old, sentences 72 months)<br>Motoric: Delayed (walked at 3 year of age)<br>Food: Problems with breast-feeding and with solid food<br>Toilet training completed by 5 years of age<br>First noted abnormalities: 6 months |
| Somatic diagnoses in infancy/childhood | Heart problem (ventricular septal defect, operated)<br>Inguinal hernia (operated)<br>Fever-induced seizures<br>Lyme disease<br>Carrier of Methicillin-resistant Staphylococcus aureus (MRSA) | Irritable bowel disorder<br>Inguinal hernia (operated)<br>Fever-induced seizures<br>Carrier of Methicillin-resistant Staphylococcus aureus (MRSA) |
| Diagnosis in research study RATSS | 299.00 Autism spectrum disorder<br>319 Intellectual disability mild<br>307.20 Unspecified tic disorder<br>307.6 Enuresis<br>307.7 Encopresis | 299.00 Autism spectrum disorder<br>319 Intellectual disability mild<br>780.52 Insomnia disorder |
| Medications at time for assessment | Levotyroxin<br>Movicol<br>Alimemazine | Levotyroxin |
| IQ screen (LEITER-R) | IQ 60 | IQ 58 |
| Adaptive functioning (ABAS 2) | GAF 40 | GAF 40 |
| Autism symptoms | ADOS-2: total score 23, severity score 9 (classified as ASD)<br>ADI-R: above cut-off (classified as ASD)<br>SRS total score: 141 | ADOS-2: total score 26, severity score 10 (classified as ASD)<br>ADI-R: above cut-off (classified as ASD)<br>SRS total score: 140 |
| ADHD symptoms | Connors 3 (raw score) | Connors 3 (raw score) |

| | | |
|---|---|---|
| | Global Index: 11<br>ADHD/Inattentive: 14<br>ADHD/Hyperactive-Impulsive: 7 | Global Index: 15<br>ADHD/Inattentive: 10<br>ADHD/Hyperactive-Impulsive: 7 |
| General comments by clinical geneticists | Abnormal eye contact<br>Monotone speech<br>Repetitive movements | Abnormal eye contact<br>Monotone speech<br>Repetitive movements<br>Echolalia |
| Morphological findings | Macrocephaly (head circumference 54 cm)<br>Wide space between teeth and late second teeth in upper jaw<br>Flat feet<br>Tapered fingers<br>Bite marks left hand | Macrocephaly (head circumference 53.8 cm)<br>Wide space between teeth and late second teeth in upper jaw<br>Flat feet<br>Tapered fingers<br>Bite marks left hand<br>Overweight |
| Parental education | Mother: University; Father: Elementary school | |

ABAS, Adaptive behavior assessment system; ADHD, Attention-deficit/hyperactivity disorder; ADI-R, Autism Diagnostic Inventory-Revised; ADOS, Autism Diagnostic Observation Schedule—Second Edition; ASD, Autism Spectrum Disorder; GAF, Global Assessment of Functioning; IQ, Intelligence quotient; SRS, Social Responsiveness Scale
